# Convergent gene expression in endometrial epithelial cells illuminates the evolution of uterine receptivity

**DOI:** 10.1101/2024.03.13.584751

**Authors:** Axelle Brulport, Małgorzata Anna Gazda, Eulalie Liorzou, Maëlle Daunesse, Bruno Raquillet, Anthony Lepelletier, Maria Sopena-Rios, Mariam Raliou, Pierrick Regnard, Lyne Fellmann, Lucie Faccin, Slaveia Garbit, Alexia Cermolacce, Ivanela Kondova, Louis Marcellin, Carole Abo, Ludivine Doridot, Camille Berthelot

**Affiliations:** Institut Pasteur, Université Paris Cité, CNRS UMR 3525, INSERM UA12, Comparative Functional Genomics group, F-75015 Paris, France; European Molecular Biology Laboratory, European Bioinformatics Institute, Wellcome Trust Genome Campus, Hinxton, Cambridge, CB10 1SD, UK; Université Paris-Saclay, UVSQ, INRAE, BREED, 78350, Jouy-en-Josas, France; SILABE – Université de Strasbourg, Fort Foch, 67207 Niederhausbergen, France; Station de Primatologie - CNRS - UAR 846, 2230 route des Quatre Tours, 13790 Rousset sur Arc, France; Biomedical Primate Research Centre (BPRC), Animal Science Department, Division of Pathology and Microbiology, Rijswijk, The Netherlands; Université de Paris Cité, Faculté de Médecine, Paris, France; Department of Gynecology Obstetrics II and Reproductive Medicine (Professor Chapron), Assistance Publique-Hôpitaux de Paris (AP-HP), Hôpital Universitaire Paris Centre (HUPC), Centre Hospitalier Universitaire (CHU) Cochin, Paris, France; Department “Development, Reproduction and Cancer”, Cochin Institute, INSERM U1016, Paris, France; Université Paris Cité, Institut Cochin, INSERM, CNRS, F-75014 PARIS, France, Paris, France

## Abstract

The epithelium of the uterine endometrium is the first maternal interface encountered by the embryo, and plays crucial roles in the maternal-embryonic crosstalk necessary to embryo implantation. Mechanisms of embryo implantation are highly variable between mammals: humans and mice have convergently evolved similar embryo implantation phenotypes, where the embryo embeds in the maternal mucosa, which differs from the ancestral mammalian and primate phenotypes. This phenomenon is thought to be partly controlled by maternal epithelial receptivity signals during the window of implantation. Here, we combined endometrial epithelial organoid models and single-cell transcriptomics to investigate how gene expression has evolved in endometrial epithelial cells between human, non-human primates and mouse at key time points in the hormonal cycle. We discovered that many maternal genes involved in uterine receptivity and embryo implantation exhibit more similar expression patterns between human and mouse compared to macaque and marmoset. In particular, we show that the endometrial expression of *LIF*, a crucial actor of endometrial receptivity in both human and mouse, is likely an evolutionary convergence rather than a conserved feature as previously hypothesised.

## Introduction

The uterus is a key organ of the mammalian female reproductive tract, with crucial roles in embryo implantation, placenta development, pregnancy support and parturition. The uterus is composed of two tissue layers: the myometrium, a thick-walled smooth muscle with supportive and contractile functions; and the endometrium, a dynamic mucosa organised into a basal and a functional layer. The functional layer experiences rapid tissue growth, then remodels to prepare for embryo implantation and eventually sheds during menstruation, while the basal layer contributes to regenerating the functional layer after menstruation (*1*). This cyclical remodelling of the endometrium is orchestrated by ovarian steroid hormones (estrogen and progesterone), which are themselves regulated by the hypothalamus-pituitary axis (*1*). The endometrium is composed of two main cell types: stromal cells, which differentiate into secretory decidual cells and participate in placenta establishment; and epithelial cells, organised into a continuous epithelial layer that lines the uterine lumen and invaginates into the stroma to line the uterine glands.

Despite its paramount importance for successful reproduction and species survival, the evolution of the mammalian uterus remains poorly understood. The tissular structures and cellular composition of the uterus are conserved across mammals, as is the overall sequence of the hormonal cycle; however, mammalian evolution is also marked by important differences in uterine morphology and physiology between species (*2*). These shifts include different mechanisms of embryo attachment and implantation into the receptive maternal endometrium (*3–5*), dramatic differences in placental invasivity into the uterine wall and in placental structures (*6*, *7*), and important variations in mechanisms of the endometrial remodelling and renewal, including several convergent acquisitions of menstruation (*8*). Capturing and comparing the molecular tenets of these time- and hormone-dependent processes *in vivo* across multiple species remains a major challenge.

Of the two main cell types that compose the endometrium, only the evolution of stromal cells has been investigated at the molecular level. Endometrial stromal cells are fibroblast cells that can differentiate into decidual cells following hormonal and embryonic cues to participate in placenta establishment (*1*). Transcriptomic comparisons of endometrial stromal cell lines across species have illuminated the functional bases of key mammalian adaptations such as decidualization (*9–11*). How endometrial epithelial cells have evolved remains however largely unexplored (*3*). The endometrial epithelium, the most external cellular layer of the endometrium, plays fundamental roles in embryo recognition and early implantation (*3*), stromal decidualization (*12*), and immunity against pathogens in the uterine cavity (*13*). As such, endometrial epithelial cells are thought to be a key player in the evolutionary changes in uterine receptivity and embryo implantation strategies observed in mammals.

Endometrial epithelial organoids are a robust model to study the functions of primary endometrial epithelial cells. Grown from tissue-derived primary epithelial cells seeded into an artificial extracellular matrix, these 3D epithelial structures recapitulate many of their *in vivo* characteristics, differentiating into secretory and ciliated cellular subtypes and secreting uterine fluid-like mucus (*14*, *15*). Here, we examine the evolution of gene expression in endometrial epithelial cells by leveraging endometrial epithelial organoids as well as single-cell transcriptomics on timed endometrial biopsies across three model mammals: human, mouse, and macaque, a non-human primate and model species for female reproductive biology. Humans and macaques diverged approximately 29 million years ago, and share many characteristics in their reproductive cycle and strategies, including similar cycle lengths (28 days), endometrial differentiation and decidualization triggered by internal maternal factors, presence of menstruations, and embryo attachment via the polar trophectoderm (*4*, *16*). Mice, which separated from primates 87 million years ago, differ significantly in their reproductive traits: menstruations are absent, and embryo attachment proceeds through different mechanisms, with recognition of the mural trophectoderm by the maternal endometrium, which triggers decidualization and endometrial remodelling (*17*). Despite significant differences in how embryo implantation proceeds between both species, human and mouse have independently evolved mechanisms of embryo implantation where the blastocyst interacts with maternal cells to become embedded into the endometrium (*3–5*, *18*). Macaque, like most primates except great apes, undergoes superficial implantation, where the embryo remains inside the uterine cavity and only superficially breaches into maternal tissues, spreading peripherally from the attachment site (*4*, *18*). These differences are thought to be at least partly driven by inter-species differences in how maternal tissues prepare to become receptive to embryo implantation in response to ovarian hormones (*17*), which is not well characterized even in model species. Here, we investigate how this first line of cellular contact between maternal and embryonic tissues has evolved between human, macaque and mouse, three widely used models of reproductive biology. We leverage a combination of endometrial epithelial organoids *in vitro* and single-cell transcriptomics experiments on tissues obtained *in vivo* to explore how transcriptomic responses to ovarian hormones have changed during evolution in the endometrial epithelium. Our results show that *LIF* and several other critical uterine receptivity genes become expressed in response to internal hormonal cues in endometrial epithelial cells in humans and mice, while these signals are absent in macaques. These signals are also absent in a more phylogenetically distant primate, the common marmoset, suggesting that they represent a convergent signature that may contribute to the phenotypic convergences in human and mouse receptivity mechanisms.

## Results

### Macaque endometrial epithelial organoids recapitulate uterine epithelium phenotypes

To investigate the evolution of the endometrial epithelium, we derived endometrial epithelial organoids from fresh endometrial biopsies of three cynomolgus macaques (*Macaca fascicularis*) and three genetically diverse mice strains (*Mus musculus*), following previously published protocols for human and mouse (**Fig. 1A, S1**; **Methods**) (*15*, *19*, *20*). We obtained endometrial epithelial organoids for both species, which continued to grow and increase in size over time, and exhibited the characteristic structure previously reported in human (*19*, *20*) and mouse (*15*) endometrial epithelial organoids (**Fig. S1**). Macaque organoids displayed the same cellular organisation as human and mouse organoids, with a polarised peripheral epithelial layer (E-Cadherin positive) surrounding a central lumen, and were proliferative (Ki-67 positive; **Fig. 1B**). These represent to our knowledge the first lines of non-human primate endometrial epithelial organoids, modelling the luminal epithelium and glands found in the macaque uterine endometrium.

**Figure 1.**
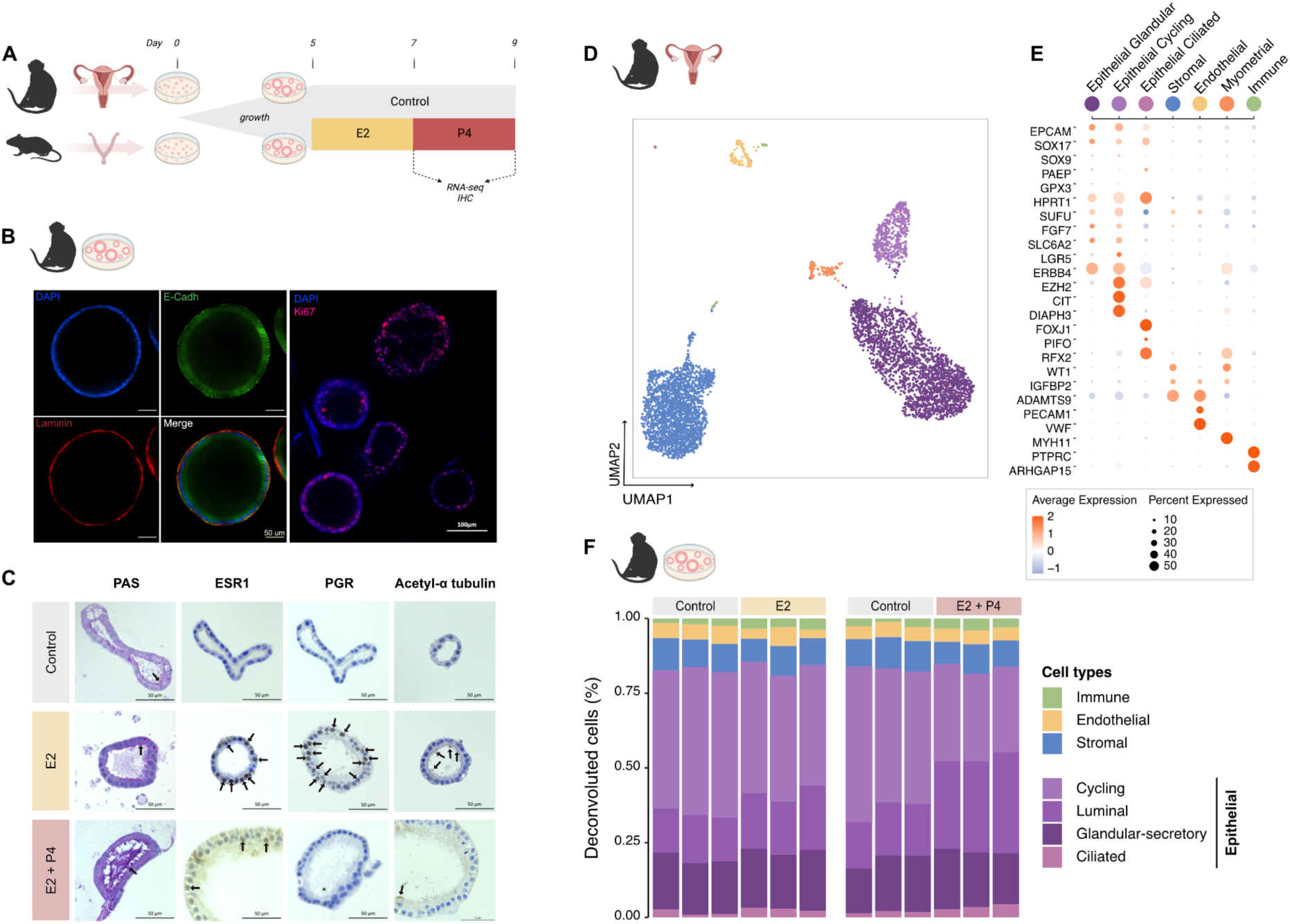
Transcriptomic landscapes of macaque hormone-responsive epithelial organoids and uterine endometrium. **(A)** Experimental design and hormonal stimulation timeline. Endometrial epithelial organoids were grown from macaque and mouse endometrial tissue samples in growth medium from day 0 to day 5, primed with E2 for 48h, then stimulated with P4 for 48h (termed E2+P4 thereafter). **(B)** Immunofluorescence staining of untreated macaque endometrial epithelial organoids for E-cadherin (epithelial cell marker), laminin (basement membrane marker) and Ki-67 (proliferation marker), x40. **(C)** Periodic Acid Schiff staining for polysaccharides (arrows), and immunohistochemistry staining for steroid hormone receptors (ESR1 and PGR; arrows) and acetylated a-tubulin to visualise cilia (arrows) in macaque organoids, x40. **(D)** Uniform Manifold Approximation and Projection (UMAP) representation of single-nuclei transcriptomes from endometrial tissue of a macaque individual in the late secretory phase. Colors correspond to the cell types in **(E)**. **(E)** Expression of representative marker genes in the different cell types identified by single-nuclei transcriptome sequencing of macaque endometrium in **(D)**. Dot size indicates the fraction of cells expressing a gene, and color indicates the scaled average expression by cell (z-score). **(F)** Cell type composition of macaque endometrial epithelial organoids estimated by cellular deconvolution of their bulk transcriptomic profiles based on published human endometrial snRNA-seq data.

We next investigated how macaque organoids respond to estrogen and progesterone, by mimicking the hormonal exposure that uterine cells experience *in vivo*. The hormonal cycle is largely conserved between mammals (*1*, *21–24*), with a first proliferative phase driven by estrogen (E2) and a second secretory phase where endometrial cells experience differentiation under the effects of progesterone (P4) (*1*). After five days of growth, we exposed macaque and mouse organoids to E2 for 48 hours, or to E2 for 48 hours followed by P4 for another 48 hours (*15*) (**Fig. 1A**). Hormonal treatment modified the phenotype of macaque and mouse endometrial organoids, which acquired a brown, dark colour, with modification of their 3D conformation and reduced growth (**Fig. S1A**). Hormone-treated organoids grew significantly more slowly than untreated controls in both species (**Fig. S1B**; macaque: p = 0.01; mouse: p = 0.01; ANOVA).

We then explored whether macaque endometrial organoids express known functional markers of hormonal response. As observed in human (*15*, *19*, *20*) and mouse (*15*), macaque organoids secrete mucus upon stimulation by estrogen and progesterone, as evidenced by Periodic Acid Schiff (PAS) staining (**Fig. 1C, S2A**). Macaque organoids also express the estrogen receptor (ESR1) after both E2 and P4 treatment, and the progesterone receptor (PGR) after E2 treatment only (**Fig. 1C**), consistent with previous histological analysis on human tissue which showed that PGR is down-regulated in the mid and late luteal phase with no expression in the functional glands (*25*). We also observed the presence of ciliated epithelial cells after E2 and P4 treatment, as evidenced by acetyl-α tubulin labelling, suggesting the presence of diverse epithelial cell types as observed *in vivo* in endometrial tissue (**Fig. 1C**). Our phenotypic observations confirm that macaque endometrial epithelial organoid lines are functional and faithfully reproduce previously described *in vivo* characteristics in response to hormonal treatment in human (*15*, *19*, *20*) and mouse (*15*).

Finally, to ascertain the functional relevance of macaque endometrial epithelial organoids, we generated single-nuclei transcriptomes of the endometrium from a macaque individual in the progesterone-controlled late secretory phase, and compared cellular composition *in vivo* to those of macaque organoids. We obtained high-quality transcriptomes from 5,520 single nuclei, and performed unsupervised cell clustering and two-dimensional embedding using UMAP for representation (**Fig. 1D-E**). Based on marker gene expression (*26*), we identified several distinct populations of epithelial cells during the secretory phase in macaque, including glandular and cycling cells as well as a small population of ciliated cells, along with other expected endometrial and uterine cell types, such as stromal, endothelial, myometrial and immune cells. Gene expression deconvolution using a comprehensive single-cell atlas of the human endometrium (*26*) estimates that macaque endometrial epithelial organoids are also largely be composed of cycling endometrial epithelial cells (control : 48%, E2: 42%, E2+P4: 30%), glandular (control : 18%, E2/E2+P4: 19%) and luminal cells (control: 16%, E2: 19% E2+P4: 31%), and small fractions of ciliated cells (control: 2%, E2/E2+P4: 3%), in line with phenotypic observations and previous reports in human endometrial organoids (*15*, *19*, *20*) (**Fig. 1F; Methods**). The estimated proportions of differentiated cells increased with hormonal stimulation (E2 vs control: p = 0.013; E2+P4 vs E2: p = 0.002; t-test). These transcriptomic comparisons confirm that macaque endometrial epithelial organoids successfully model the endometrial epithelium *in situ* and are an appropriate system to study the behaviour of these cells. Similarly, we confirmed that mouse endometrial epithelial organoids recapitulate previously reported phenotypes (*15*) and display a cellular composition consistent with mouse endometrium, in particular recapitulating the absence of ciliated cells in mice compared to humans and macaques (**Fig. S2**). The successful establishment of functional macaque endometrial epithelial organoids reported here therefore presents an opportunity to investigate the evolutionary dynamics of the uterine epithelium transcriptome in mammals.

### Transcriptomic responses to hormones reveal conserved functional pathways in the endometrial epithelium between mammals

To understand how endometrial organoids respond to hormones across mammalian models for reproductive biology, we compared the transcriptomes of hormone-treated and control organoids from macaque and mouse with two publicly available RNA-seq datasets from human endometrial epithelial organoids in similar hormone stimulation experiments (**Methods**). Principal component analysis (PCA) revealed that transcriptomic profiles clearly distinguish between control and hormone-treated organoids in all three species (**Fig. S3**). We identified 630, 1,346 and 2,520 differentially expressed genes (DEGs) after E2 treatment and 4,472, 3,228 and 2,322 after E2+P4 treatment in human, macaque and mouse, respectively. (**Fig. S4; Methods**). Examining the function of the top DEGs across species revealed key actors of endometrial receptivity (*PGR, ESR1, Asgr2, RPLP2*) (*27–30*), embryo implantation (*KLK11, TPPP3, CDC20B, Gper1*) (*27*, *28*, *31–33*), decidualization (*TPPP3, Gper1, GJA1, PLXNA4, C1rb, Ceacam10*) (*32*, *34–41*), pregnancy outcomes (*GSTA3, Kcnj1*) (*35*, *42*, *43*) and endometrial dynamic changes during the menstrual cycle (*ESR1, PGR, DEUP1, LGALS3*) (*44–46*). Additionally, in each species, around 75% of genes differentially expressed under E2 treatment were also DEGs after E2+P4 treatment (**Fig. S4, S5**), suggesting that we capture a continuous response of endometrial epithelial organoids to E2 and E2+P4 hormonal exposure. We note, however, that E2+P4 treatment elicited a stronger transcriptomic response than E2 alone in both primates, with many genes becoming expressed only in the presence of progesterone, while the difference between both treatments was more commensurate in mouse.

To compare gene expression between human, macaque and mouse, we identified 15,015 one-to-one orthologous genes across the three species (**Methods**). Transcriptome comparison supports that the identity of endometrial epithelial cells is largely conserved between primates and mice, with a generally high correlation of their gene expression levels (Spearman correlation; human-macaque: p = 0.75; human-mouse: p = 0.71; macaque-mouse: p = 0.80; **Fig. 2A**) and as previously observed in other tissues and cell types (*47*, *48*). PCA revealed that the two first principal components (PC) separate the transcriptomes by species, and together explained 82.3% of the total variance in gene expression (**Fig. 2B**). Gene expression changes induced by hormonal treatment in each species were minor compared to the divergence in gene expression accumulated over evolutionary time. PC3 correlates with treatments in human and macaque samples and explains 5.2% of the total gene expression variance (**Fig. S6A**), while mouse samples separated with treatment along PC5 (1.6% of total variance; **Fig. S6B**), suggesting that primate endometrial epithelial cells share common features in their response to E2 and P4 that are not conserved in mouse cells.

**Figure 2.**
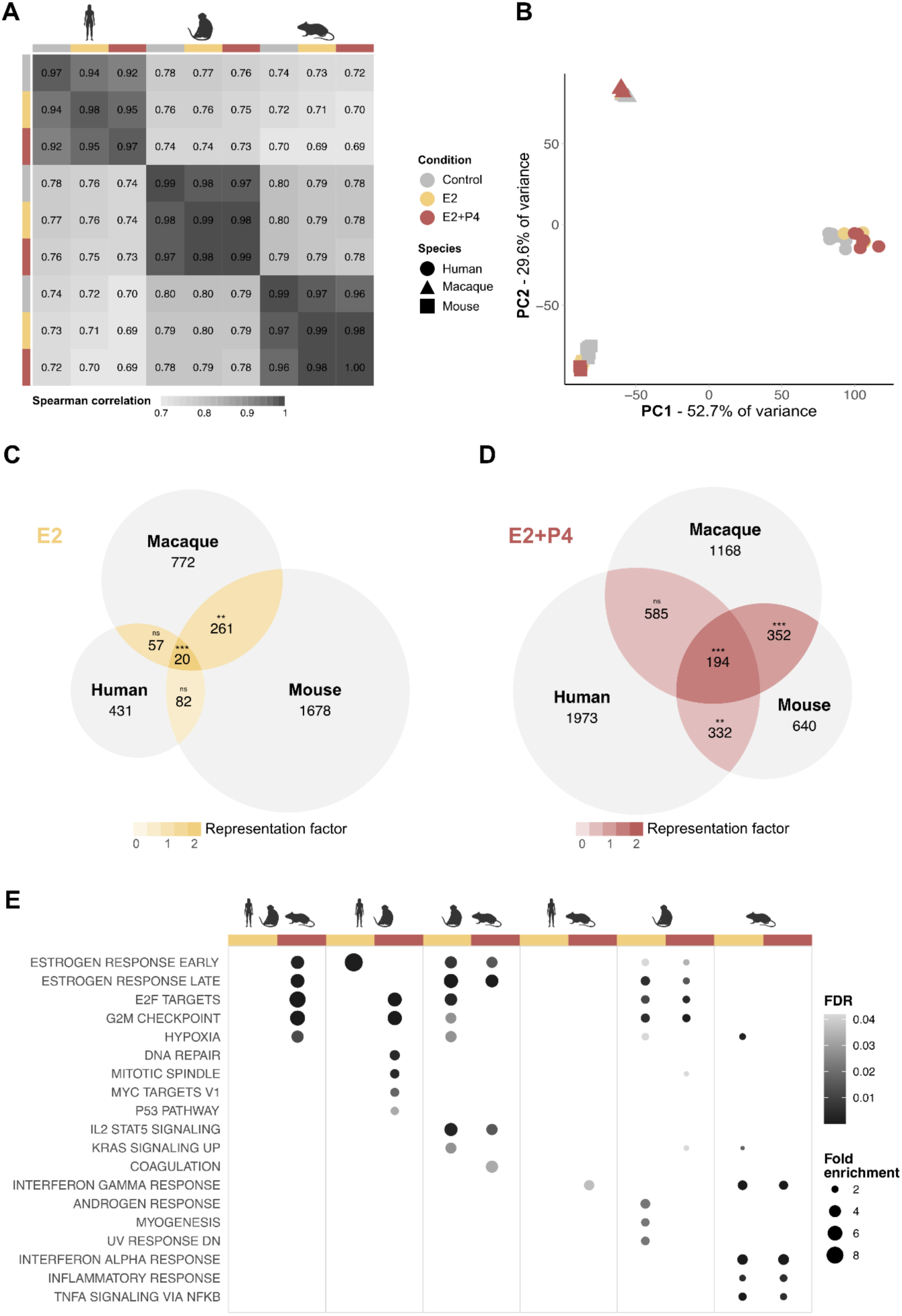
Endometrial epithelial cell transcriptomes are overall conserved between model mammals, but display divergences in hormonal response. **(A)** Gene expression correlations across conditions and species. Each coefficient represents the mean Spearman rho coefficient across all pairwise correlations of biological replicates per condition and species. **(B)** Principal component analysis of gene expression across 15,015 one-to-one orthologues between all three species. **(C-D)** Differentially expressed genes shared between human, macaque and mouse **(C)** after E2 treatment, and **(D)** after E2+P4 treatment. Colour intensity represents the representation factor, e.g. observed to expected overlap. *: p-values < 0.05, **: p-values < 0.01 and ***: p-values < 0.001, hypergeometric test, BH adjustment. **(E)** Hallmark gene sets annotations over-represented in genes differentially expressed in all species, in any pair of species, or only in one species in response to hormonal treatment. P-values were calculated using a hypergeometric test followed by false discovery rate (FDR) correction.

We then investigated whether hormonal treatment induces changes in gene expression shared across the three species. We identified a core set of differentially expressed genes common to all species after each treatment (E2: 20 genes; E2+P4: 194 genes; **Fig. 2C-D, Table S1**), as well as subsets of DEGs shared by any pair of species. Functional enrichment analysis using Hallmark gene sets (*49*) revealed that genes responding to hormones and shared between species are enriched in pathways involved in cell growth (E2F target, G2M checkpoint, DNA repair, mitotic spindle, KRAS signalling) but also and more interestingly, in functions with key roles for maternal preparation to embryo implantation (**Fig. 2E, Table S2**; **Methods**). These include estrogen response, hypoxia pathways (*50*), the p53 pathway (*51*, *52*), inflammation (interferon gamma response)(*53*, *54*) and immune signalling pathways (IL2 STAT5 signalling) (*55*, *56*). Amongst those shared DEGs, we identified several proposed biomarkers for endometrial receptivity such as *GADD45A* and *MAOA* (*57*). Species-specific DEGs were also enriched in inflammatory and immune response genes. Altogether, and in all three species, hormonal stimulation modifies a range of functional pathways with described roles in embryonic implantation, consistent with the role of endometrial epithelial cells as the first line of contact with the implanting blastocyst in mammals. Embryonic implantation however proceeds differently across our three study species. Human and mouse have convergently evolved modes of embryo implantation where the blastocyst breaches through the endometrial epithelium and becomes embedded into the maternal mucosa (*3–5*). Macaque on the other hand displays superficial embryo implantation, where the blastocyst remains at the endometrium surface before placentation proceeds, which is the ancestral phenotype of primates (*4*, *5*). In the next paragraphs, we take advantage of this experimental design to explore in more detail how ovarian hormones prime maternal epithelial cells in species that have evolved different strategies of embryonic implantation.

### Uterine receptivity genes are convergently activated in human and mouse endometrial epithelial organoids

Uterine receptivity and endometrial preparation to embryo implantation are poorly annotated in functional databases, which limits our ability to dissect how implantation preparation pathways have evolved in endometrial cells. To circumvent this, we curated a list of 59 genes with well-described roles in uterine receptivity, embryo attachment and embryo implantation in mammalian endometrial epithelial cells from the literature, termed “Implantation”, which we added as an additional process to the analysis (**Table S3**; **Methods**).

To further explore whether cellular processes are similarly activated or repressed in response to hormonal stimulation across human, macaque and mouse, we extracted all genes annotated to the functions highlighted in **Fig. 2E**, regardless of whether they are detected as differentially expressed. We then tested whether expression fold-changes of these genes in response to hormonal stimulation were correlated across species. Human and macaque organoids displayed stronger correlations in gene expression fold-changes for most processes, consistent with their closer phylogenetic distance (**Fig. 3A**). Cell cycle processes showed high correlations between all conditions except in human organoids after E2 treatment, where they were anti-correlated to other samples. This may reflect a difference in cell expansion dynamics of human epithelial cells in response to estrogen specifically, or could also reflect a technical effect, as these samples were processed externally to our lab. Pathways related to hormone response showed higher correlation within primates (**Fig. 3A**), in line with our previous observation that both human and macaque cells exhibit a common gene expression response overall to hormonal treatment (**Fig. S6A**). In contrast, genes involved in maternal preparation to implantation and the p53 pathway showed significant correlations in expression fold-changes between human and mouse, which were not shared with macaque (**Fig. 3A**). Interestingly, the p53 pathway has a well documented role during implantation in both human (*52*) and mouse (*51*), and controls the endometrial expression of the Leukemia Inhibitory Factor (*LIF*) (*51*), an interleukin family 6 cytokine pivotal to maternal-embryonic crosstalk and early implantation (*58*, *59*).

**Figure 3.**
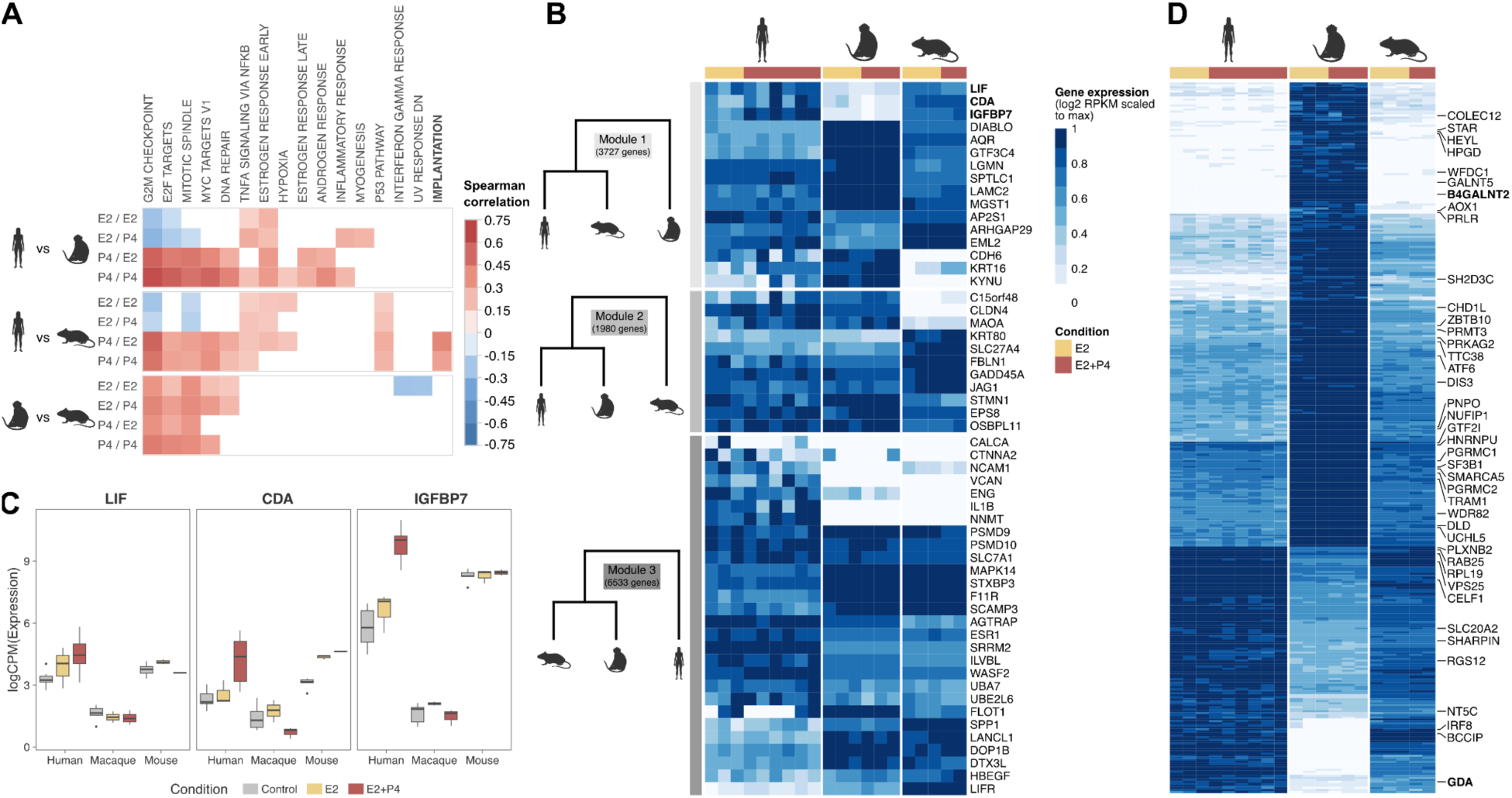
Maternally-expressed genes related to uterine receptivity share more similar expression patterns in human and mouse compared to macaque. **(A)** Pairwise correlations of expression log2-fold-changes in response to E2 or E2+P4 treatment compared to controls, for all genes annotated to over-represented Hallmark gene sets, as well as maternally-expressed genes with documented roles in endometrial receptivity and embryo implantation (termed ‘Implantation’). Only correlations with a BH-adjusted p-value < 0.1 are represented. **(B)** Expression of maternal implantation genes across treatments in human, macaque and mouse endometrial epithelial organoids (normalised RPKM, log2-transformed, scaled to the maximum expression value for each gene). Genes are divided in three groups, corresponding to the main modules obtained in the co-expression analysis (**Methods**), which reflect differences in phylogenetic similarity as shown on the left. **(C)** Normalised expression levels for *LIF*, *CDA* and *IGFBP7* across treatments and species. **(D)** Normalised expression levels for 352 genes strongly co-expressed with *LIF* (co-expression weight > 0.4). Labelled genes have documented roles in endometrial receptivity, decidualization, embryo implantation, pregnancy outcomes and ovarian hormone response. Log2-transformed expression levels are scaled to the maximum value for each gene for representation.

Because genes involved in uterine receptivity and maternal preparation for implantation display an unusual pattern of expression convergence between human and mouse, we next sought to identify additional candidates by exhaustively searching for similar convergent expression patterns. We performed a Weighted Gene Correlation Network Analysis (WGCNA, **Methods**) which identified 8 modules of co-expressed genes, where the three largest modules reflect stronger expression similarities between either human and macaque, human and mouse, or macaque and mouse (**Fig. 3B, S7**). Module 1 corresponds to a group of 3,727 genes more similar in expression between human and mouse than with macaque. Sixteen genes involved in implantation are in Module 1, and in particular *LIF*, *CDA* and *IGFBP7*, which are expressed in human and mouse epithelial organoids but exhibit low expression in macaque (**Fig. 3C**). *LIF* is essential to successful pregnancy establishment, and is generally thought to be a deeply conserved actor of embryo implantation in mammals due to its importance for uterine receptivity in both human and mouse (*58–60*). The low expression of *LIF* in macaque epithelial cells after hormonal stimulation was therefore striking, and suggests a potentially significant difference in maternally-controlled mechanisms of endometrial receptivity and preparation to embryo implantation. *CDA* and *IGFBP7* have been overall less described, but also have documented roles in endometrial receptivity (*61–64*). To identify other candidate genes of potential interest within Module 1, we prioritised 352 genes most strongly co-expressed with *LIF* (weight > 0.4; **Table S4**). Half of those genes (178) encode membrane-bound proteins, and many of these candidate genes have been previously described as involved in endometrial decidualization and implantation (*HEYL, VPS25, RGS12, GDA, SHARPIN, EIFAEBP1, RPL19, IRF8, HPGD, SF3B1, DIS3, GTF2I, SMARCA5, PGRMC1, COLEC12, PRLR, GALNT5, B4GALNT2, PRMT3, CELF1, SH2D3C, ATF6, UCHL5, GNA12*) and pregnancy outcomes (*PLXNB2, RAB25, TRAM1, SLC20A2, PRLR, CHD1L, WDR82, TTC38, DLD*, *PRKAG2, BCCIP, AGGFI, NUFIP1*), or are regulated by estrogen and/or progesterone (*PNPO, NT5C, RAB25, HNRNPU, WFDC1, AOX1, STAR, ZBTB10, PGRMC2*) (**Fig. 3C**).

These analyses revealed that a large panel of known actors in endometrial receptivity and maternal preparation to embryo implantation, notably *LIF*, present more similar expression and hormone-triggered variations between human and mouse than between human and macaque. These differences in gene expression are therefore functional candidates that may have contributed to the convergent similarities of embryo implantation mechanisms in human and mouse, which both differ from macaque.

### Single-cell transcriptomics of the endometrium confirms a convergent evolution of epithelial gene modules in human and mouse

While our previous analyses highlight candidate genes that may have experienced convergent expression changes in human and mouse, other interpretations may also explain those results: first and foremost, endometrial organoids may not entirely faithfully replicate gene expression *in vivo*. Second, these differences in gene expression may be macaque-specific traits, unshared with other primates that also exhibit superficial implantation. To address those potential caveats, in addition to our transcriptomic atlas of the endometrium in macaque, we also performed single-nuclei transcriptome sequencing of the endometrium in a common marmoset and in mice, collected in non-fecundated female individuals at the time when the endometrium becomes receptive to embryo implantation (late secretory phase of the uterine cycle in marmoset; day 3.5 of pseudopregnancy in mouse; **Methods**). Marmoset is a platyrrhine (New World monkey) that also presents a superficial embryonic implantation phenotype, which is the likely ancestral phenotype in primates (*65*). Additionally, we used the single-nuclei transcriptomes of human endometrial samples in the mid and late secretory phase available from the Reproductive Cell Atlas (*26*).

We profiled the transcriptomes of a total of 35,368 high-quality nuclei (9,297 in marmoset and 20,551 in mouse, in addition to the 5,520 in macaque described in **Fig. 1**), and identified endometrial epithelial cell populations based on documented marker expression and comparison to the human data (**Fig. 4A**; **Fig. S8**; **Methods**). We confirmed that some human endometrial epithelial cells display expression of *LIF* during the secretory phase (**Fig. 4B**), mainly in glandular cells where 8% of nuclei have detectable *LIF* expression in this dataset (average normalised expression: 26 CPM; **Fig. 4D**). In the mouse pseudopregnant endometrium, *LIF* is also expressed by glandular epithelial cells at relatively high levels, with 20% of nuclei detected as *LIF*-positive across the glandular cell cluster at the beginning of the window of implantation in this species (average normalised expression: 156 CPM; **Fig. 4A-B**).

**Figure 4.**
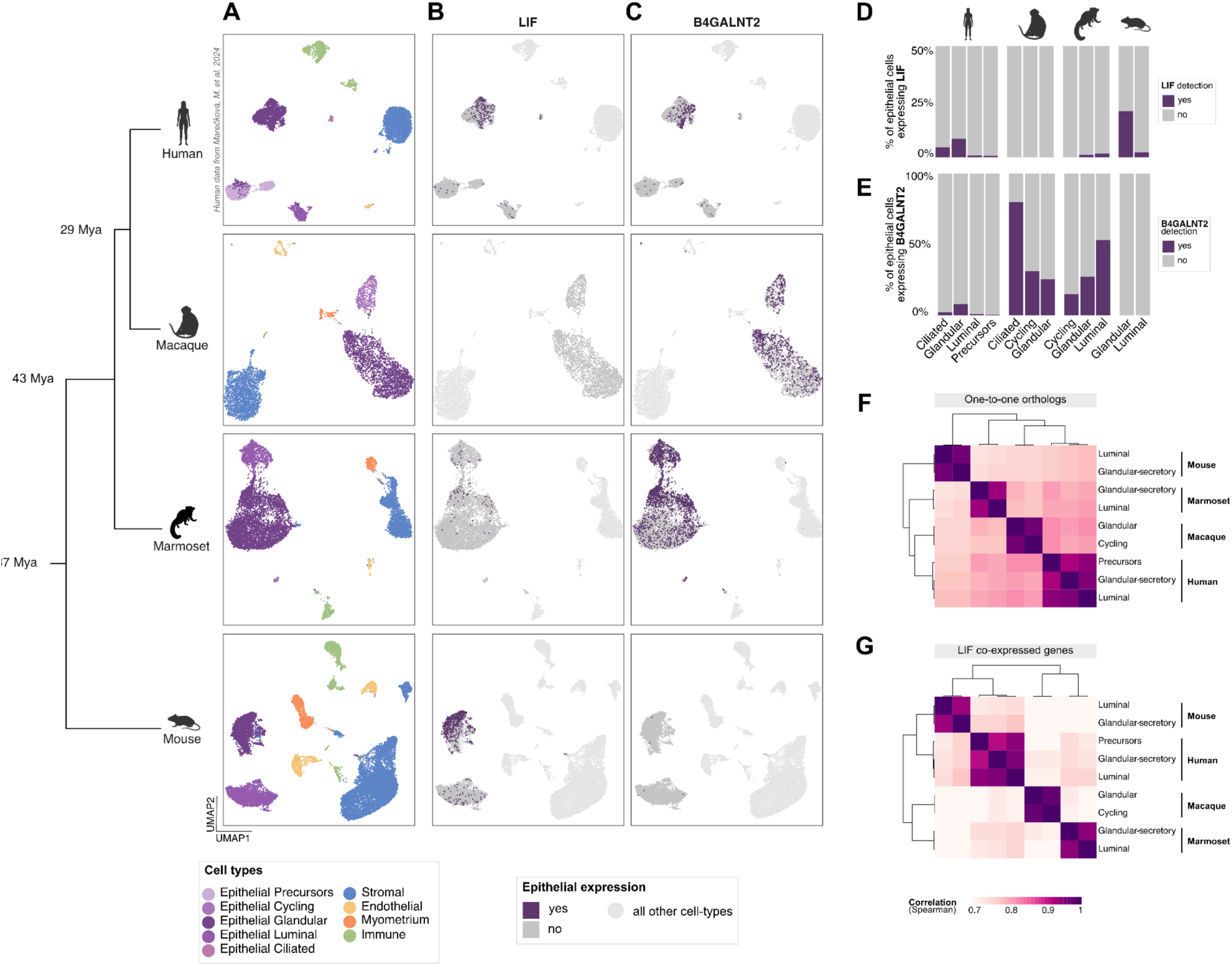
Single-nuclei transcriptomics of the endometrium confirms the convergent evolution of LIF and other co-expressed maternal genes during the window of implantation in human and mouse in vivo. **(A)** Uniform manifold approximation and projection (UMAP) of single-nuclei transcriptomes of late secretory phase endometrium from human, macaque and marmoset, and pseudopregnant mouse at 3.5 DPC, corresponding to the window of implantation in this species. Cell types were annotated using representative marker genes as in Figure 1D and Figure S7. **(B-C)** Expression of *LIF* **(B)** and *B4GALNT2* **(C)** in epithelial cells in each species. Epithelial cells where expression was detected (gene count ≥ 1) are colored in purple. Epithelial cells where no expression was detected are colored in grey. Other cell types are colored in light grey. **(D-E)** Fractions of cells expressing *LIF* **(D)** and *B4GALNT2* **(E)** across epithelial cell subtypes in each species. **(F-G)** Correlation of gene expression levels across epithelial cell subpopulations (with ≥ 100 cells) in each species for **(F)** all one-to-one orthologs and **(G)** 232 one-to-one orthologous genes strongly co-expressed with *LIF* in organoids (out of 352 candidates; 120 genes did not have a one-to-one ortholog in marmoset). Hierarchical clustering was performed based on Spearman correlations.

In contrast, *LIF* expression was not detected at all in rhesus macaque cells in secretory phase, and only in rare cells not specific to a given epithelial cell type in marmoset (**Fig. 4A-B; Methods**). These results further support that *LIF*-mediated signalling in the secretory phase has changed during primate evolution, and is not triggered by maternal ovarian hormones alone in either a New World or an Old World monkey. We also investigated the expression of *CDA* and *IGFBP7*, but found that these genes are expressed in marmoset epithelial cells similarly to human and mouse, suggesting that their low expression in macaque cells is specific to that species or clade (**Fig. S9**). However, we highlight *B4GALNT2* as another example of a gene of potential interest with different expression patterns in human and mouse versus macaque and marmoset, both in organoids and *in vivo* (**Fig. 4C,E**). *B4GALNT2* mediates the catalysis of Sd(a), an important blood antigen also expressed at the surface of epithelial cells and recognized by the DBA lectin. *B4GALNT2* has previously been implicated as an essential actor of embryo recognition and attachment (*66*, *67*), and can be expressed either maternally or embryonically in different mammalian species (*67–69*), suggesting that modifications in the localization of the Sd(a) antigen may play a role in mammalian differences in embryo implantation mechanisms, which to the best of our knowledge has not been explored.

Finally, we investigated whether candidate genes co-expressed with *LIF* in organoids, identified in **Fig. 3D**, also display convergent expression in human and mouse *in vivo*. We measured the correlation of gene expression levels in endometrial epithelial cells across all four species. When considering all orthologous genes, transcriptomes group as expected based on phylogenetic proximity, and human epithelial cells are more closely correlated to macaque and marmoset cells (**Fig. 4F**). However, clustering based on the expression of genes co-expressed with *LIF* groups human and mouse epithelial cells together, and apart from macaque and marmoset (**Fig. 4G**). This analysis *in vivo* therefore confirms that a network of hundreds of co-expressed genes, including *LIF*, have experienced coordinated changes in endometrial epithelial cells resulting in more similar expression in human and mouse compared to other non-human primates. As these genes include many known actors of decidualization, receptivity and embryo implantation, this convergent transcriptomic signature during the window of implantation represents the first evidence for a molecular mechanism that may explain the similarities in implantation phenotypes between human and mouse, paving the way for future functional investigations.

## Discussion

We report here the first evolutionary comparison of endometrial epithelial cell transcriptomes across three major models of female reproductive biology - human, macaque and mouse - and their modifications under ovarian hormone stimulation. Cyclic gene expression changes are essential to the functions of the uterine endometrium, which have been profoundly remodeled along mammalian evolution (*3*, *4*, *9*, *17*, *70*). This rapid evolution of a crucial reproductive interface may be driven by an ongoing evolutionary crosstalk between mother and embryo, which co-evolve to optimize and negotiate their respective fitnesses (*71*). In that context, the evolutionary dynamics of the uterine epithelium are of particular interest as the frontline of interactions between maternal and embryonic tissues. While the cellular identities of endometrial epithelia are conserved between primates and mouse, as revealed by the overall conservation of their transcriptomic profiles, our results highlight that their response to hormones has changed over time. Across all three species, and consistent with the proliferation of endometrial cells observed in all mammals along the hormonal cycle, ovarian hormones activate or repress a core network of genes involved in cell cycle regulation and steroid hormone response. In all three species, pathways modified in epithelial cells first by estrogen and then by progesterone were functionally related, suggesting a continuous tuning of epithelial functions along the cycle. However, we observed a marked divergence in epithelial cell response to hormones in each species, with both primate species sharing more similarities than with the more distantly related mouse. While hormones modified gene expression in pathways with either described or suspected roles in uterine receptivity and embryo implantation across all three species, these transcriptomic changes displayed low overlaps and correlations between species, providing a molecular background for their divergent reproductive phenotypes.

The processes of embryo recognition and implantation are highly variable across mammalian species, illustrated by the varying degrees to which the embryo will invade into maternal tissues during early implantation between different species and the orientation of the blastocyst relative to the maternal mucosa (*4*, *17*, *18*). While these differences are well documented at the phenotypic level, the molecular mechanisms that mediate differences in embryo implantation are largely unknown, especially where they imply maternal-controlled processes. In this study, we identify an unexpected correlation between human and mouse in the expression of maternal genes with well-supported roles in uterine receptivity and embryo recognition, attachment and implantation, suggesting that some of the independently-acquired similarities in their implantation phenotypes coincide with convergence at the gene expression level. Our analyses reveal in particular that *LIF*, a key actor of the maternal-embryonic cross-talk, shows differences in expression during the window of implantation between human, non-human primates, and mouse. *LIF* has long been recognized as a main trigger of maternal receptivity and embryo recognition in the mammalian uterus (*72*, *73*), and *LIF* is an important predictor of fertility and receptivity in humans, where the gene is expressed in epithelial endometrial cells throughout the secretory phase, peaking during the implantation window (*74*, *75*). A recent study in mouse uteri detailed how maternal expression of *Lif* in glandular epithelial cells initiates epithelial crypt formation and stromal differentiation for embryo embedding in the maternal mucosa (*60*), confirming the direct relevance of this gene to the mechanistic process of embryo invasion in the endometrium. Strikingly, our results reveal that *LIF* is not expressed during the secretory phase in the absence of fecundation in macaques and marmosets, where embryo implantation remains substantially more superficial than in either human or mouse. We note that *LIF* is also essential to successful implantation in primates with superficial implantation, including macaques (*76*): our results however suggest that differences in the timing and regulation of *LIF* and other receptivity genes may play a role in the ability of the endometrium to support deep invasion by the embryo. One intriguing hypothesis could be that in primates with superficial implantation, increase of maternal *LIF* expression might be triggered by embryonic signals rather than internal hormonal cues alone, possibly resulting in a more superficial receptivity phenotype in the endometrium. Exploring other candidates highlights that many membrane-localized proteins and known actors of maternal-embryo recognition present similar correlations of expression between human and mouse, but not macaque or marmoset, expanding the hypothesis that evolutionary tinkering in this early crosstalk may be important to implantation phenotypes. Interestingly, another recent study investigating peri-implantation endometrial gene expression in guinea pig and opossum also reported inter-species variations in the expression of *LIF* and other receptivity genes, further supporting that rapid divergence of maternal signalling is a key feature of early maternal-fetal interactions in placental mammals (*77*).

One major roadblock to elucidating the evolution of uterine processes in mammals is the robust functional annotation of female reproductive pathways, which lags behind that of other more widely studied organs. Trophectoderm and placental genes have been more thoroughly investigated in humans and other species (*78–82*), in part because these tissues are more accessible either naturally at parturition or through in-vitro fecundation and blastoid models. Early maternal contributions to these processes remain obscure due to the difficulty of capturing these timed events *in vivo* or modeling them *in vitro*, and therefore are not integrated in functional databases, resulting in a form of self-perpetuating ascertainment bias. Our results highlight how this annotation void can mask functional differences in critical pathways, as manual curation of genes with well-supported roles in uterine receptivity was decisive to uncover molecular convergence between human and mouse gene expression programs.

Overall, our results illustrate how inter-species comparisons combining *in vitro* and *in vivo* transcriptomics approaches can identify differences in the expression of functionally relevant genes and pathways to understand the molecular tenets of phenotypic change and convergent evolution. This approach is especially important for rapidly evolving traits, such as reproductive phenotypes, where comparisons between pairs of model species such as human and mouse may not be sufficient to capture functionally relevant change. We anticipate that mechanistic studies dissecting how the molecular actors identified here interact and control maternal-foetal communication will further illuminate how endometrial epithelial cells have evolved to modulate embryonic implantation in mammals.

## Materials and Methods

### Macaque tissue sampling

Macaque experiments were approved by the CEEA35 Ethical Committee (ethical approval number APAFIS#19437-2019022515049585) for two individuals and by the Paris-Saclay Ethical Committee (ethical approval number APAFIS#20525-2019050616506478) for the third one. *Macaca fascicularis* females aged between 8 and 13 years were used for this study. For two animals, endometrial biopsies were collected with a Novak curette (1 or 2 mm; GCCI medical: 70-5902-01, 70-5902-02) by vaginal route under ultrasound control and under general anaesthesia. One was menstruating (day 2 of the menstrual cycle) and the other was in the luteal phase (day 25 of the menstrual cycle). For the third individual, half of the uterus was opportunistically collected during the euthanasia of a control animal in an unrelated study. Cycle phase was unknown for this animal. The uterine mucosa was separated from the myometrium by scalpel scraping. For all three animals, endometrial fragments were preserved in tissue storage solution (MACS Tissue Storage Solution, Miltenyi Biotec, 130-100-008) with 1% antibiotics (Penicillin/Streptomycin, Gibco, 15140-148) at 4°C for 24 hours prior to generation of the organoid lines.

### Mouse tissue sampling

Mice experiments were approved by the Pasteur Institute Ethical Committee (dha220002 – EvoMens). Female mice aged between 10 and 32 weeks were used for this study. We used genetically diverse mice from the Collaborative Cross (CC012, CC042 and CC002 strains, one individual each) to approximate the genetic diversity of our other two study species, which are outbred. Mice were kept in the Pasteur Institute animal housing facility at constant temperature, humidity and day/night cycle, with access to water and food *ad libitum*. Mice were euthanized by cervical dislocation during estrus, detected by vaginal smears tests, and the whole uterus was used for organoid generation. Organoid cultures were immediately established.

### Endometrial epithelial organoid cultures

All cell culture steps were identical between macaque and mouse organoids, except for minor differences in growth media composition as previously published between human and mouse (*15*, *19*, *20*) (**Table S5, S6**).

Macaque endometrial biopsies or mouse uterine horns were rinsed with DPBS without Ca2+/Mg2+ (Gibco, 14190144), and cut into small pieces with dissecting scissors in a Petri dish containing 200 ul of ice-cold DMEMF12 (Gibco, 12634-010) on ice. Tissue samples were transferred to a C-tube (Miltenyi Biotec, 130-093-237) and both enzymatically and mechanically dissociated in 8 ml of Liver Digest Medium (Gibco, 17703034) containing 1% antibiotics (Penicillin/Streptomycin, Gibco, 15140-148) using a GentleMacs tissue dissociator (Miltenyi Biotec, 130-096-427) on manufacturer’s program Multi_B01 at 37°C. Tissue dissociation was completed by syringe trituration, passing the solution through 18G and 21G needles, 10 times each. Digestion was stopped with ice-cold DMEMF12 and the supernatant was sequentially filtered through a 100 µm cell strainer (Falcon, 352360) and a 30 µm cell strainer (Miltenyi Biotec, 130-098-458). The fraction retained above the 30 µm filter, containing mainly undigested or partially digested glands, was recovered. Both fractions (cells and glands larger than 30 um, and cells smaller than 30 µm including digested glands) were centrifuged 10 min at 300g at 4°C. Both pellets were resuspended in ice-cold 80% Matrigel (Corning, Growth Factor Reduced, 356231)/20% DMEMF12 (Gibco, 12634-010) at 1:20 ratio (vol/vol), and 25 µl droplets were plated in pre-warmed 48-well plates (Corning, 3548). After solidification, 250 µl of growth medium were added in each well (**Table S5** for growth media composition for mouse, and **Table S6** for macaque). Medium was refreshed every 2 to 3 days. Organoid cultures were passaged every 8 to 12 days by manual pipetting at a ratio of 1:3 to 1:5 depending on growth efficiency. From the first passage, the cultures from the fractions above 30 µm and below 30 µm were pooled. Organoids of low passage number (P1 – P3) were used for all experiments.

### Organoid hormonal treatment

Mouse and macaque endometrial epithelial organoids were treated for two consecutive days with 17-ß estradiol (E2) (Sigma Aldrich, E8875) at 10 nM or for two days with 17-ß estradiol (E2) (Sigma Aldrich, E8875) at the same concentration following by two days with progesterone (P4) (Sigma Aldrich, P8783) at 1µM. At the end of each experiment, endometrial organoids were removed from Matrigel with 250 µl of Cell Recovery Solution (Corning, 354253) per well (1h at 4°C) and centrifuged at 400g for 10 min at 4°C. Fixations for immunochemistry and immunofluorescence were performed immediately, as detailed below. Before RNA extraction, the organoids were cryopreserved in Recovery™-Cell Culture Freezing Medium (Gibco, 12648010) and stored at −80°C in a Mr. Frosty™ freezing container (Thermo Fisher, 5100-0001) for 24 hours and then in liquid nitrogen.

### Immunofluorescence and confocal microscopy

Organoids were fixed for 30 min in formalin (Roth, P087.5) at room temperature, centrifuged and the organoid pellet resuspended in 70% ethanol. Organoids were permeabilized using PBSFT (FBS 10% (Gibco, 10270-106) and Triton X100 0,5% (Sigma Aldrich, 9036-19-5) in DPBS (Gibco, 14190144)), which was also used as incubating and rinsing solvent during the immunostaining process. Primary antibodies were incubated overnight at 4°C (**Table S7** for a detailed list of antibodies). Organoids were centrifuged at 300g for 10 min at 4°C and washed twice with PBSFT. Secondary antibodies were incubated overnight at 4°C (all from Novus Biologicals: DyLight 488 donkey anti-mouse IgG (NBP1-75125) or DyLight 650 donkey anti-rabbit IgG (NBP1-75644) at 1:200 and DAPI (Invitrogen, R73606)). Final rinsing was done with DPBS (Gibco, 14190144) and immunostained organoids were placed in a 96-well flat bottom plate (Corning, 3474), in 100 µl of DBPS (Gibco, 14190144). Immunofluorescence microscopy was performed on a Nikon eclipse Ti2, CSU-W1 confocal microscope, using x20 or x40 lenses and NIS elements AR, software v.5.11.02.

### Immunohistochemistry and Periodic Acid Schiff staining

After removing the growth medium, organoids were removed from the Matrigel as above, embedded in 4% agarose (Sigma, A9639), fixed overnight in formalin at 4°C, dehydrated and paraffin wax-embedded. Sections of 3 µm were subjected to immunohistochemical staining for ESR1, PGR and acetyl-α tubulin (**Table S7** for antibodies used). Antigen retrieval was performed with a citrate buffer at pH6 for PGR and acetyl-α tubulin, and Tris EDTA buffer at pH9 for ESR1, at 100°C for 20 min. This was followed by detection with streptavidin-HRP-conjugated secondary antibodies (LEICA, Bond Intense R Detection System, DS9263). Sections were counterstained with haematoxylin (Sigma, GHS2128) for 5 min, dehydrated and mounted. Negative controls were performed using mouse and rabbit sera (1/1000, Jackson ImmunoResearch, 015-000-120 and 011-000-120) as recommended by The Histochemical Society’s Standards of Practice for Validation of Immunohistochemical Assays (*83*). Periodic Acid Schiff (PAS) staining was performed to visualise mucin. Paraffin sections were dewaxed, rehydrated and immersed in 1% periodic acid (Sigma, P7875) for 7 min, rinsed in water, treated in Schiff’s reagent (Sigma, 1.09033.0500) for 20 min, rinsed in water, counterstained with Harris haematoxylin (Sigma, GHS2128) for 1 min, washed, immersed in a saturated solution of lithium carbonate (Sigma, 62470-500G-F), washed, dehydrated and mounted. Pictures were taken using a ZEISS Zen.Apotome.2-upright widefield microscope at x40 magnification and Zen blue 2012 imaging software.

### Organoid size measurements

Organoid size was assessed by measuring the largest axis. For each condition, measurements were made at day 5, 7 and 9 on all organoids present in the same plane of two images taken randomly in two different wells. Pictures were taken using an Olympus CKX53 microscope at x10 magnification and analysed with ImageJ software. Organoid sizes were compared using 1-way analysis of variance (ANOVA). This experiment was performed across 3 biological replicates for each condition.

### RNA extraction and quality assessment

This experiment was performed as triplicates, with the exception of the E2+P4 experiment in mouse, where only two duplicates were successfully extracted. RNA was extracted from organoids using the RNeasy® Plus Micro Kit with on column DNAse treatment (Qiagen, 74034) according to the manufacturer’s instructions. RNA quality and quantity were assessed using Agilent RNA nano or pico chips on a Agilent BioAnalyzer 2100 and a Qubit with Qubit RNA HS Assay Kit (Thermo Fischer, Q32852), respectively (**Table S8**).

### Bulk RNA-seq library preparation and sequencing

Library preparations were prepared using the Illumina® Stranded mRNA prep, ligation kit (Illumina, 20040532) according to the manufacturer’s instruction from 35 ng of RNA (except for macaque replicate_2, for which 5 ng were used for all experimental conditions). Library concentrations were assessed using a Qubit instrument with dsDNA HS Assay (Thermo Fischer, Q32854), and library fragment sizes were assessed using Agilent High Sensitivity DNA chips on an Agilent BioAnalyzer 2100 instrument. Equimolar pools of 8 libraries including one replicate of each condition and species were sequenced with NextSeq 1000/2000 P2 reagents 100 cycles kit using an Illumina NextSeq2000 instrument. Libraries were sequenced as single-end 108 bp reads with an average library size of 62 millions reads (**Table S8**).

### Human organoid data curation

We curated publicly available data from two equivalent experiments in human organoids, which were stimulated with E2 and P4 over a similar time-course (*26*, *84*). The first dataset was sequenced as bulk transcriptomes using Illumina paired-end reads, while the second was sequenced as single cells using 10XGenomics Chromium GEX library preparation and Illumina sequencing. Both of those works included both biological and technical replicates, and we only extracted one technical replicate per biological sample. In total, the human dataset consists of 7 controls, 3 E2 treatments and 7 E2+P4 treatments. Of note, some E2+P4 samples were also treated with cAMP, a stromal decidualization factor; however, transcriptomic analyses revealed that those samples grouped with other E2+P4 samples and that cAMP treatment had little impact on the transcriptome, so we elected to retain those as well.

### Transcriptome analysis of organoid experiments

#### Reference transcriptomes

The reference transcriptomes for human (GRCh38.p13), crab-eating macaque (Macaca_fascicularis_6.0), and mouse (GRCm39) were downloaded from Ensembl 108 (*85*).

#### Orthologous gene selection

To perform cross-species comparisons, we extracted one-to-one orthologous protein coding genes present in all three species (human, mouse and crab-eating macaque) using Ensembl Biomart 108, resulting in 15,015 genes.

#### Bulk RNA sequencing analysis

Quality of bulk RNA-seq libraries, produced for this study and from published data, was controlled using FastQC v.0.11.9. Low quality bases and adapters were removed using Trimmomatic v.039 (*86*). Trimmed reads were pseudo-mapped using Salmon v.1.9.0 (*87*) to the appropriate reference transcriptome. Quality control statistics were extracted from the report generated by Salmon during mapping. Transcript quantifications were converted into gene quantifications using tximport v.1.24.0 (*88*).

#### Reanalysis of published human organoid single-cell transcriptomes

For single-cell RNA-seq data from human organoids, we followed the standard mapping procedure of 10xGenomics CellRanger v.6.0.1 (*89*) to the reference human transcriptome. The output was processed in Seurat v.4.2.0 (*90*). Cells were retained if they met the following quality criteria: a minimum of 250 genes were detected (nFeature_RNA), a minimum of 500 unique molecules detected (nCount_RNA), a log10 of number of genes per UMI (log10GenesPerUMI) higher than 0.80, and fewer than 20% of mitochondrial reads. After removing low quality cells, gene expression was aggregated and extracted as pseudo-bulk quantifications (PseudobulkExpression).

#### Batch effect removal

Gene expression quantification files were combined into count tables per species. Macaque and mouse libraries were corrected for technical batch effects due to sequencing batches using CombatSeq (sva v3.44.0 R package) (**Fig S3 A-B**) (*91*). Human libraries were corrected for technical batch effects due to sample origin.

#### Differential expression

Differential expression analysis was performed using DeApp v.1.7.4 (*92*). Only genes with expression levels over 1 CPM in at least two samples were considered as expressed and retained for analysis. DeApp performs differential expression using DeSeq2 v.1.38.2 (*93*), edgeR v.3.38.4 (*94*) and limma v.3.52.4 (*95*). Genes were considered as differentially expressed if they had an adjusted p-value below 0.05 for all three methods. Volcano plots shown in **Fig. S4** were constructed using the statistics obtained with limma. We note that statistics obtained across all three methods were highly consistent (Spearman correlations between log fold-changes ≥ 0.95) and method choice did not alter the general conclusions.

#### Cell type deconvolution

Cell type deconvolution of endometrial epithelial organoids bulk RNAseq was performed using Dampened weighted least squares with default parameters (DWLS v0.1.0, (*96*)). Because the endometrial epithelial organoids derived *in vitro* were exposed to E2 or E2+P4 treatment to mimic *in vivo* hormonal exposure of the endometrium, we expect epithelial organoids to model epithelial cells from the entire hormonal cycle. Therefore, deconvolution was performed using the human single-nuclei atlas from Marečková et al. 2024 (*97*) as reference, which includes samples from healthy donors spanning both proliferative and secretory phases. Samples from donors under hormonal treatment were excluded, as well as the MUC5B epithelial cell cluster likely originating from cervical epithelial mucosa as proposed by the authors of the dataset (*97*).

### Cross-species analysis

#### Transcriptome correlation

Expression counts for the 15,015 one-to-one orthologs were normalised across species using edgeR size factors. Transcriptome correlation was performed on all one-to-one orthologs using log2-transformed normalised expression counts. Mean Spearman correlation coefficients across conditions and species were represented using the R packages stats v.4.2.2 and pheatmap v.1.0.12. Principal component analysis (PCA) was performed on log2-transformed and z-normalised expression counts using R package FactoMineR v.2.6.

#### Cross-species differentially expressed genes

We compared changes in gene expression in response to hormones across species. For each species, DEGs were restricted to one-to-one orthologous genes and intersected across species for each treatment independently. We tested the significance of the intersection across species by performing a hypergeometric test with Benjamini-Hochberg (B-H) p-value adjustment.

#### Functional enrichment

Functional enrichment analysis was performed with ShinyGO v.0.77 (*98*) using the Human Hallmark gene set annotations (*49*). We used sets of shared DEGs for each combination of species as foreground to perform functional enrichment analysis. Background genes included all 13,008 orthologous genes expressed more than 1 CPM in at least two samples for each species (e.g. all genes that have been tested for differential expression in all three species).

#### Curation of maternal implantation gene set

We manually curated a list of maternally expressed genes with described roles in endometrial receptivity or embryo implantation from recent reviews (*61*, *99*) and research articles on implantation mechanisms (**Table S3)**. Genes were included when they were expressed in endometrial epithelial cells during the window of implantation and if they were experimentally associated with implantation and endometrial receptivity in at least two independent studies (**Table S3**).

#### Gene expression fold-change correlations

Expression fold-change values between treatments and controls were calculated with limma for all one-to-one orthologous genes in the over-represented Hallmark gene sets and in the maternal implantation gene set. Spearman correlations of log-fold-change values were calculated for each gene set between species and conditions. Correlation p-values were adjusted for multiple testing using Benjamini-Hochberg correction. Because we considered a fairly large number of pathways, each containing a relatively small number of genes, we used a relaxed significance threshold for this analysis (adjusted p-value < 0.1).

#### Gene co-expression analysis

Co-expression analysis across species and conditions was performed on the one-to-one orthologs genes using the WGCNA v.1.71 (*100*) R package. The weighted gene co-expression network was constructed using as input edgeR factor-scaled log2 RPKM expression levels for the three species in the treated samples and a soft thresholding power (β) of 6, resulting in 13 co-expression gene modules (**Fig. S7**). Genes with a co-expression weight with *LIF* higher than 0.4 were considered as candidates of interest for further analysis.

### Single-nuclei RNA-seq

#### Tissue collection

A half snap-frozen uterus in mid-luteal phase from a female rhesus macaque individual (*Macaca mulatta*) was sourced from the Biomedical Primate Research Center (BPRC) biobank. All procedures regarding the maintenance of the Tissue bank at BPRC were in accordance with the Dutch law on animals, and the European legislation concerning the care and use of laboratory animals. The BPRC is accredited by the American Association for Accreditation of Laboratory Animal Care (AAALAC) International and is compliant with the European directive 2010/63/EU as well as the “Standard for Humane Care and Use of Laboratory Animals by Foreign Institutions” (National Institutes of Health, ID A5539-01). Cycle phase was confirmed histologically by an experienced veterinary pathologist. Thin cryostat organ sections were obtained and the endometrial part of the tissue was identified by hematoxylin eosin staining and dissected on serial sections with a cold scalpel before processing.

A whole uterus in mid-luteal phase was obtained from a female common marmoset individual (*Callithrix jacchus*) after approval from the CEEA71 Ethical Committee (ethical approval number APAFIS#31395-20210325103447). Cycle phase was determined by plasma progesterone titration by Enzyme Linked Fluorescent Assay (MiniVIDAS® Progesterone 30409, Biomérieux, France) over 15 days prior to surgery, and confirmed histologically after surgery. Hysterectomy was performed under general anaesthesia when progesterone levels plateaued to the maximum levels observed in this species (139,43 ng/uL). The uterus was immediately rinsed with cold PBS 1x and dissected on ice to separate the endometrium from the myometrium. Endometrium samples were snap-frozen in liquid nitrogen and stored at −80°C before processing.

For pseudopregnant mice, 3 females from the B6CBAF1/J mouse strain were mated with a vasectomised male, and pseudopregnancy was timed at 0.5 day if a vaginal plug was assessed in the morning post mating. At 3.5 days post *coitum* (DPC), whole uteri were collected and were immediately rinsed with cold PBS 1x, snap-frozen in liquid nitrogen and stored at −80°C before processing. In mice, sterile mating induces a neuroendocrine response leading to a pseudopregnancy status characterised by a prolactin surge and the increased production of progesterone during 6-8 days (*101*, *102*).

#### Nuclei isolation

For the rhesus macaque sample, all buffers were prepared in agreement with demonstrated protocol CG000375_RevC from 10xGenomics for Complex Tissues for Single Cell Multiome ATAC + Gene Expression Sequencing. A few adaptations were made to the nuclei isolation protocol. Briefly, the endometrial tissue fragments were incubated for 5 min in 2 ml of lysis buffer and homogenised 25 times using a dounce on ice before adding 2 ml of wash buffer. The suspension was filtered through a 100 um filter and centrifuged for 5 min at 500g and 4°C. The pellet was washed a second time with 1 ml of wash buffer. After centrifugation, nuclei were resuspended in 500 µl of wash buffer and labelled with an anti-nuclear pore complex proteins antibody (clone Mab414) and sorted on a FACSAria instrument (BD Biosciences), with a 100 nozzle on a cold block. The collection tube was then cold centrifuged for 7 min at 500g and the nuclei pellet was incubated for 2 min on ice with lysis buffer containing digitonin to permeabilize the membranes. After a final washing step with 1 ml of wash buffer and centrifugation, nuclei were resuspended in 49 µl of diluted nuclei buffer to obtain 12,500 nuclei in 5 µl of buffer for further processing.

For both mouse and marmoset, endometrial samples were embedded in OCT and thinly sliced with a scalpel on ice. The fragments were transferred to a dounce, incubated for 5 min in 1 ml lysis buffer (10mM Tris-HCL pH7.5, 146 mM NaCl, 1 mM CaCl2, 21 mM MgCl2, 0.03% Tween-20, 0.01% BSA, 1 mM DTT, 10% EZ Lysis Buffer (Sigma/Merck), 0.5 U/µl RNase Inhibitor) and then ground with a pestle. The homogenate was then filtered through a 70 um strainer, rinsed with 400 µl of wash buffer (250 mM sucrose, 10mM Tris-HCL pH7.5, 5 mM MgCl2, 1% BSA, 1 mM DTT, 0.5 U/µl RNase Inhibitor) and centrifuged for 5 min at 500g and 4°C. The pellet was washed a second time and centrifuged again. The nuclei pellet was finally resuspended in 200 ul of wash buffer and filtered through a 40 µm filter. 15,000 nuclei were loaded in the 10XGenomics Chromium instrument for each species.

#### 10X Genomics Chromium library preparation and sequencing

For rhesus macaque, nuclei were proceeded immediately with Chromium Next GEM Single Cell Multiome ATAC + Gene Expression Reagent Bundle Kits according to the Chromium Next GEM Single Cell Multiome ATAC + Gene Expression User Guide (CG000338_RevF). For mouse and marmoset, nuclei were proceeded immediately with Chromium Next GEM Single Cell 3ʹ Kit v3.1 according to Chromium Next GEM Single Cell 3ʹ Reagent Kits v3.1 (Dual Index) User Guide (CG000315_RevE). Equimolar pools of 4 libraries including one replicate of each species were sequenced with NextSeq 1000/2000 P2 reagents 100 cycles kit using an Illumina NextSeq2000 instrument. Libraries were sequenced as paired-end following the instructions specific to each 10xGenomics chemistry (Multiome or Gene Expression). Equimolar pools including one replicate of each condition and species were sequenced using an Illumina NextSeq2000 instrument.

#### Raw reads processing

For macaque, mouse and marmoset, CellRanger v.7.1.0 was used for demultiplexing with the corresponding Chromium chemistry. CellRanger transcriptome files were built using *mkref* function and with the default parameters for each species using the respective genome assemblies (rheMac10, mm10, calJac4). CellRanger count function was used for the generation of the count matrix using the intronic mode (--include-introns) and default parameters to map reads to the transcriptome and generate a count matrix for every UMI. CellRanger prefiltered count matrices (filtered_feature_bc_matrix) without empty droplets were used for downstream analysis.

#### Nuclei quality filters and data processing

Quality control was performed for each sample independently using Seurat v.5.0.1 (*103*). SoupX v.1.6.2 (*104*) correction was applied following standard recommendations to correct the count matrix for ambient RNA contamination. Nuclei expressing fewer than 500 genes, more than 5000 genes and/or more than 5% of mitochondrial reads after SoupX correction were filtered out as low quality. Doublet identification and removal was performed using DoubletFinder v 2.0.4 (*105*) following standard guidelines (pK auto-estimated, pN=0.25, N=15000 corresponding to the expected number of nuclei loaded during library preparation). For the macaque, additional filtering was performed after data processing to remove a cluster of cells expressing both epithelial and stromal markers that were likely remaining doublets, particularly as most doublets estimated from DoubletFinder were removed from that cluster. Normalisation, feature selection, scaling and dimension reduction was performed using standard procedures with Seurat (*see* **Table S7** for snRNA-seq QC and processing metrics). Clustering was then performed using the Louvain algorithm and following Seurat guidelines (*90*, *103*). Epithelial cell clusters were identified using expression of canonical epithelial markers (*EPCAM, SOX17*), and further refined for the glandular, luminal, ciliated and proliferating epithelial clusters using marker genes from the Human Reproductive Cell Atlas (*97*) and analysis of the top marker genes by cluster (FindAllMarkers with min.pct=0.25, logfc.threshold=0.1; **Table S10-11-12** for top marker genes by cluster per species). Other broad endometrial cell types were also identified across all species (**Fig. 4A, S8**). Ciliated epithelial cells were not confidently identified in marmoset, possibly due to the number of cells as ciliated cells are rare; a small fraction of luminal epithelial cells were expressing *FOXJ1* gene, a known marker of ciliated cells, suggesting that a small population of ciliated cells may have been included in this cluster.

Finally, depth-normalised pseudo-bulk expression levels were extracted for all epithelial clusters (AggregateExpression). Epithelial clusters expression levels are presented as log_2_-transformed CPM for the one-to-one orthologous genes of interest. Pseudo-bulk performed on cell populations of less than 100 cells were not considered for transcriptome correlation analysis in **Fig. 4F-G**, resulting in the removal of macaque ciliated cells (n=10), marmoset cycling cells (n=73) and human ciliated cells (n=92).

#### Human single-nuclei dataset curation

Human endometrial single-nuclei RNA-seq data was obtained pre-processed and annotated from the Human Reproductive Cell Atlas (*97*). Only samples corresponding to healthy donors, mid-secretory and mid-late-secretory phases were retained to match the time point in the other species. Additionally, cell types annotated as MUC5B were removed as those cells likely originate from cervical epithelium as described by the authors (*97*). After subsetting the single-nuclei atlas, samples matching our inclusion criteria were re-processed following standard Seurat guidelines, and integrated using Harmony integration from Seurat v5 (*103*). The human cell type annotation was derived from the author’s existing annotation.

### Licences

Human, macaque and mouse icons were extracted from PhyloPic (http://phylopic.org), and are available for reuse under the Public Domain Mark 1.0 license.

## Acknowledgements

We thank C.M. Fovet and F. Relouzat (CEA, Fontenay-aux-Roses, FR) for providing serendipitously collected tissue samples; H. Kaessmann (ZMBH, Heidelberg, DE), C. Dégletagne (CRCL, Lyon, FR) and R. Vento-Tormo (Wellcome Trust Sanger Institute, Hinxton, UK) for sharing protocols and technical advice with tissue collection and single-nuclei sequencing; S. Chouzenoux (Institut Cochin, Paris, FR) for technical advice with cell culture; B. Ouwerling and T. Haaksma (BPRC, Rijswijk, NL) for technical support with tissue collection. We thank the scBiomarkers UTechS platform, C2RT, and the Histopathology core facility (Institut Pasteur, France), for support in conducting this study. We thank O. Godefroy, A. Dujardin and A. Coudray of the BIME EOPS facility and G. Pinto Moreira, J. Gourlay and C. Daviaud of the PEAC facility of C2RA for providing pseudopregnant mice tissues (Institut Pasteur, France). We thank M. Monot and the Biomics platform, C2RT, Institut Pasteur, Paris, France, supported by France Génomique (ANR-10-INBS-09) and IBISA. We thank M. Andrieu and V. Karunanithy from the Cytometry and Immunobiology Facility (CYBIO) and F. Letourneur from the GENOM’IC facility, Institut Cochin, Paris, France, for their help with single-nuclei processing.

## Author’s contributions

A.B. and C.B. designed experiments; A.B., B.R. and A.L. performed experiments with help from M.R. and L.D.; M.A.G., E.L., M.D., and M.S.R. analysed the data; L.F., P.R., P.M.V., L.F., A.C., S.G., L.M. and C.A. performed animal tissue collections; I.K. provided tissue samples and histopathology expertise; E.L., M.D., A.B., M.A.G. and C.B. designed figures; C.B., A.B., E.L. and M.D. wrote the manuscript with input from M.A.G and other authors.

## Data availability

All sequencing data generated in this study has been deposited to the NCBI Gene Expression Omnibus (GEO; https://www.ncbi.nlm.nih.gov/geo/) under accession numbers GSE260471 (bulkRNA-seq: GSE260470; snRNA-seq macaque and marmoset: GSE260468) and GSE286078 for pseudopregnant mouse snRNA-seq. All original code for this study is accessible on GitLab at https://gitlab.pasteur.fr/cofugeno/endometrium_organoid_2024 under the GNU Lesser General Public License (LGPL).

## Supplementary Figures

**Figure S1.**
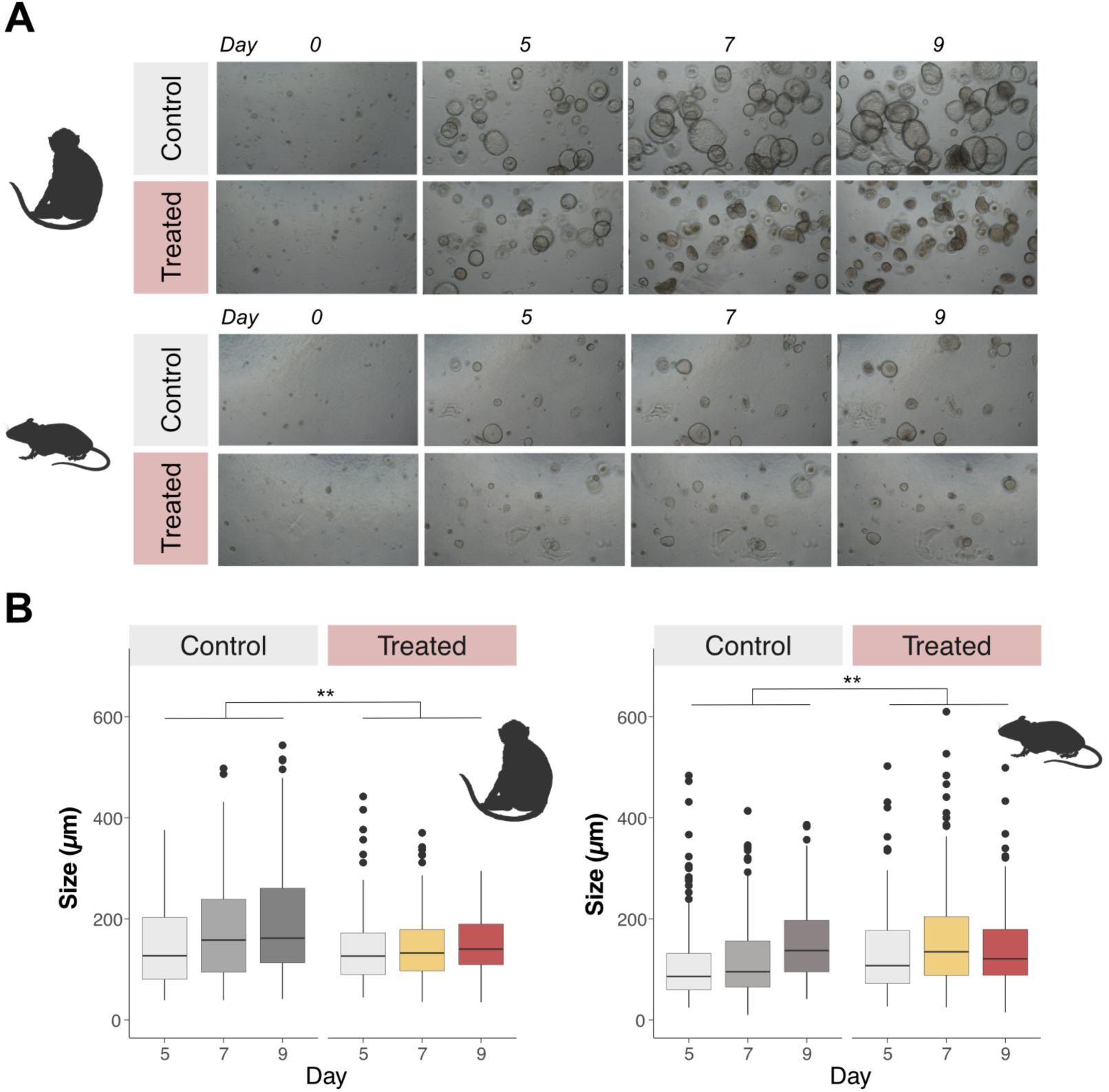
Growth assessment of macaque and mouse organoids. **(A)** Growth of control and treated endometrial epithelial organoids monitored over time for both species after seeding (day 0), after growth step (day 5), after E2 treatment (day 7) and after P4 treatment (day 9), x10. **(B)** Diameters of macaque and mouse organoids before treatment (day 5), after E2 treatment (day 7) and after E2+P4 treatment (day 9). **: p-values < 0.01, ANOVA.

**Figure S2.**
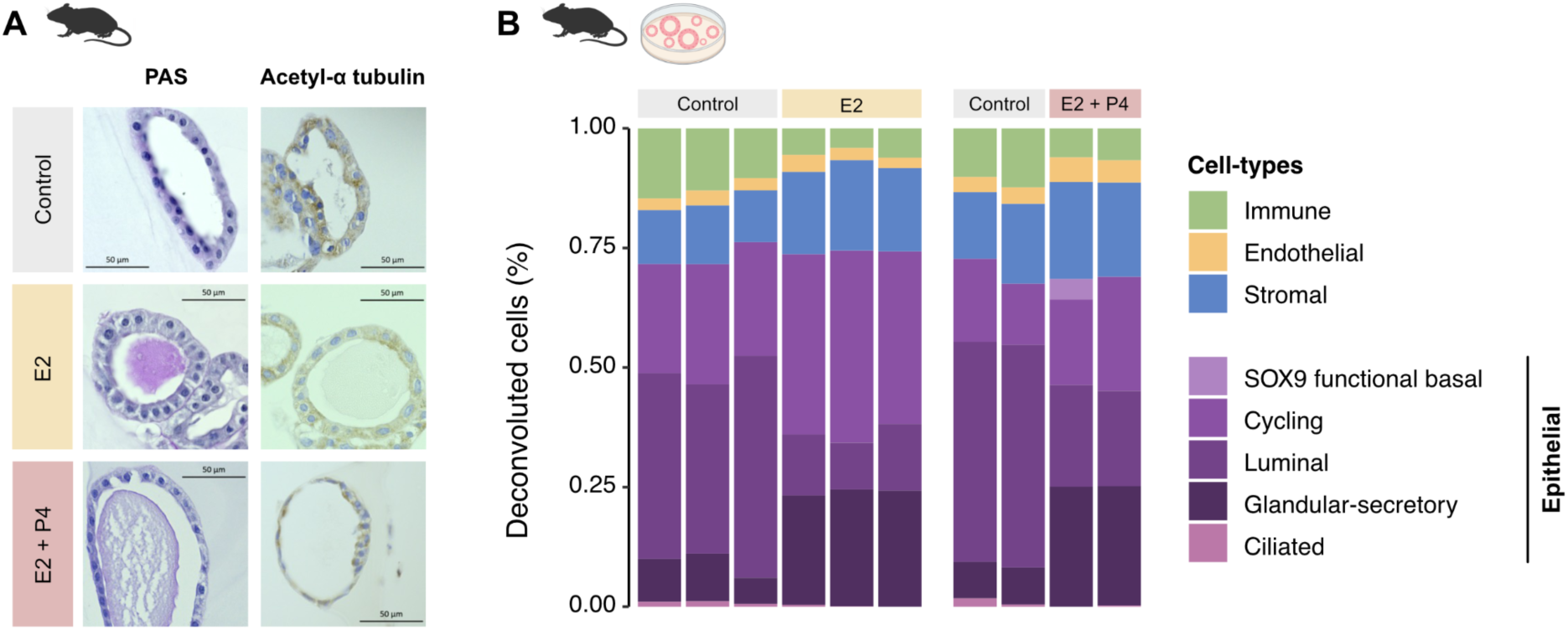
Histological and transcriptional characterisation of mouse endometrial epithelial organoids. **(A)** Periodic Acid Schiff (PAS) staining for polysaccharides; immunohistochemistry staining for acetylated alpha-tubulin, confirming the absence of ciliated cells in mouse organoids, x40. **(B)** Cell type composition of mouse endometrial epithelial organoids estimated by cellular deconvolution of their bulk transcriptomic profiles based on published human endometrial snRNA-seq data.

**Figure S3.**
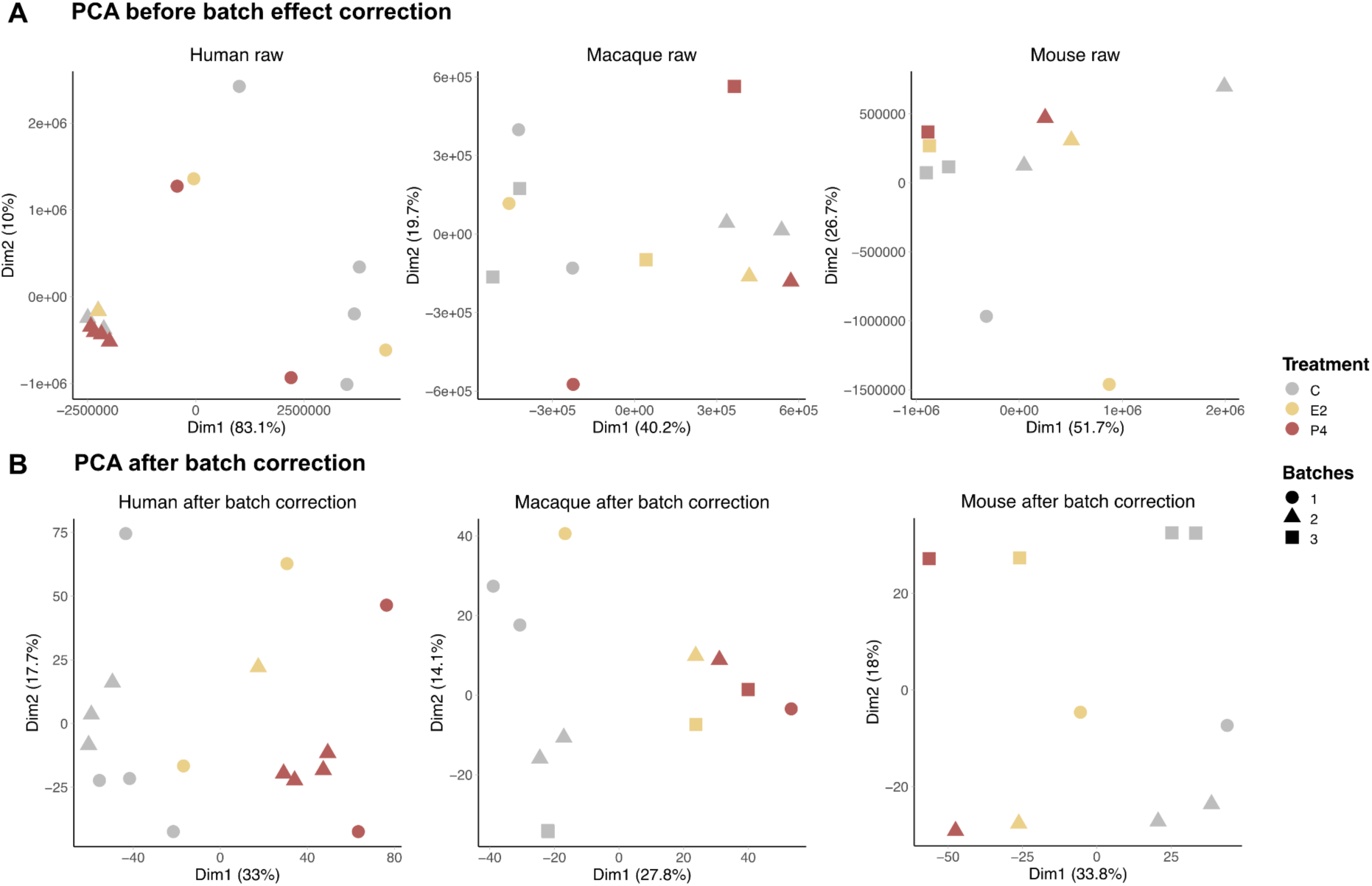
PCA of bulk RNA-seq from human, macaque and mouse organoids. **(A)** with raw counts (**B**) and counts after batch correction. Samples are coloured by condition. Counts were z-normalised and scaled before PCA.

**Figure S4.**
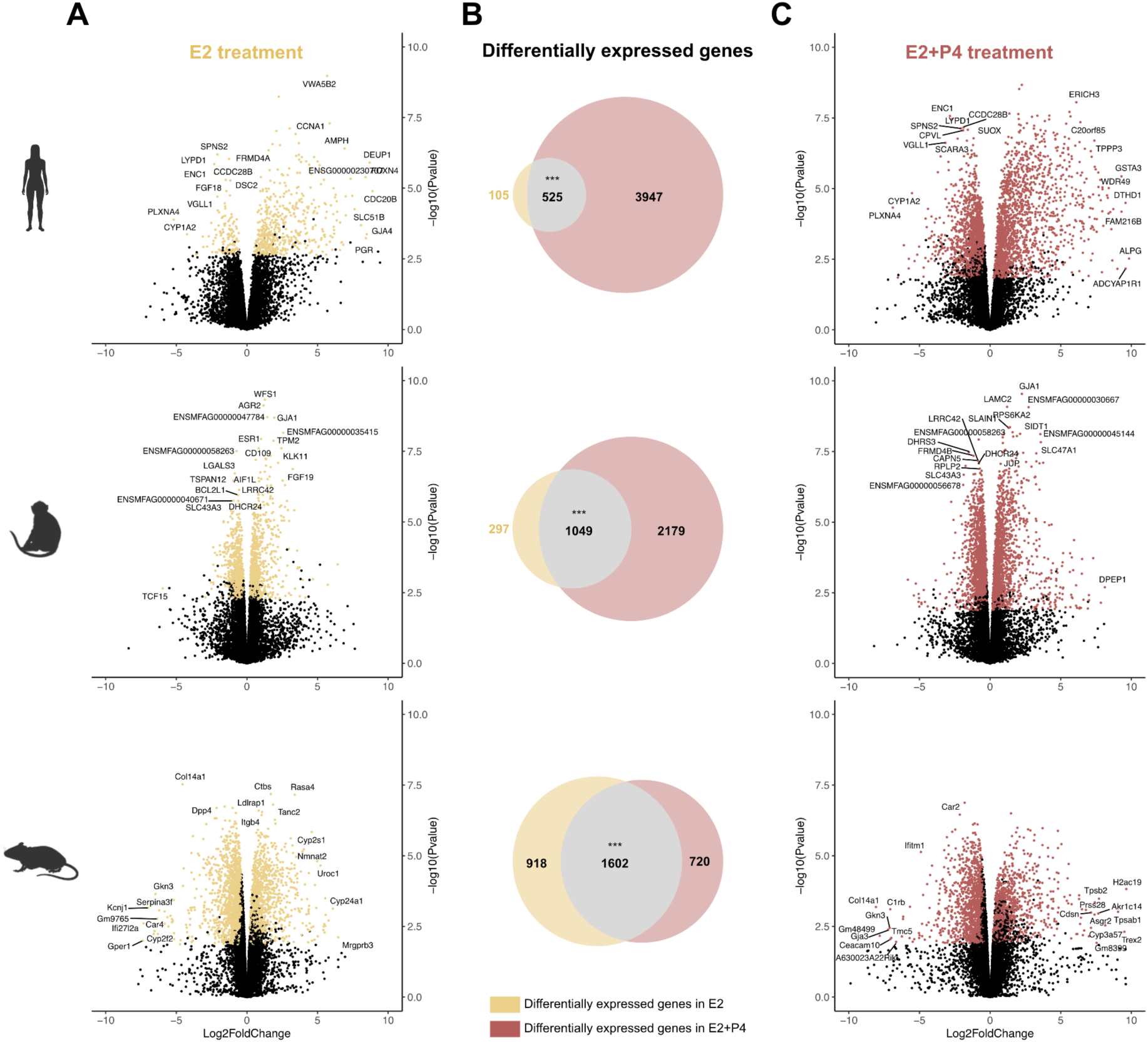
Differential expression analysis in human, macaque and mouse organoids after E2 or E2+P4 treatments. **(A,C)** Volcano plots of differential gene expression in response to E2 (yellow, **A**) or E2+P4 (red, **C**) treatment for each species. Gene names are reported for the top 10 up- and down-regulated DEG based on highest Euclidean distance to 0. **(B)** Venn diagram of the overlapping DEGs between E2 and E2+P4 conditions, colour intensity represents the representation factor for each species. ***: p-values < 0.001, hypergeometric test, BH adjustment.

**Figure S5.**
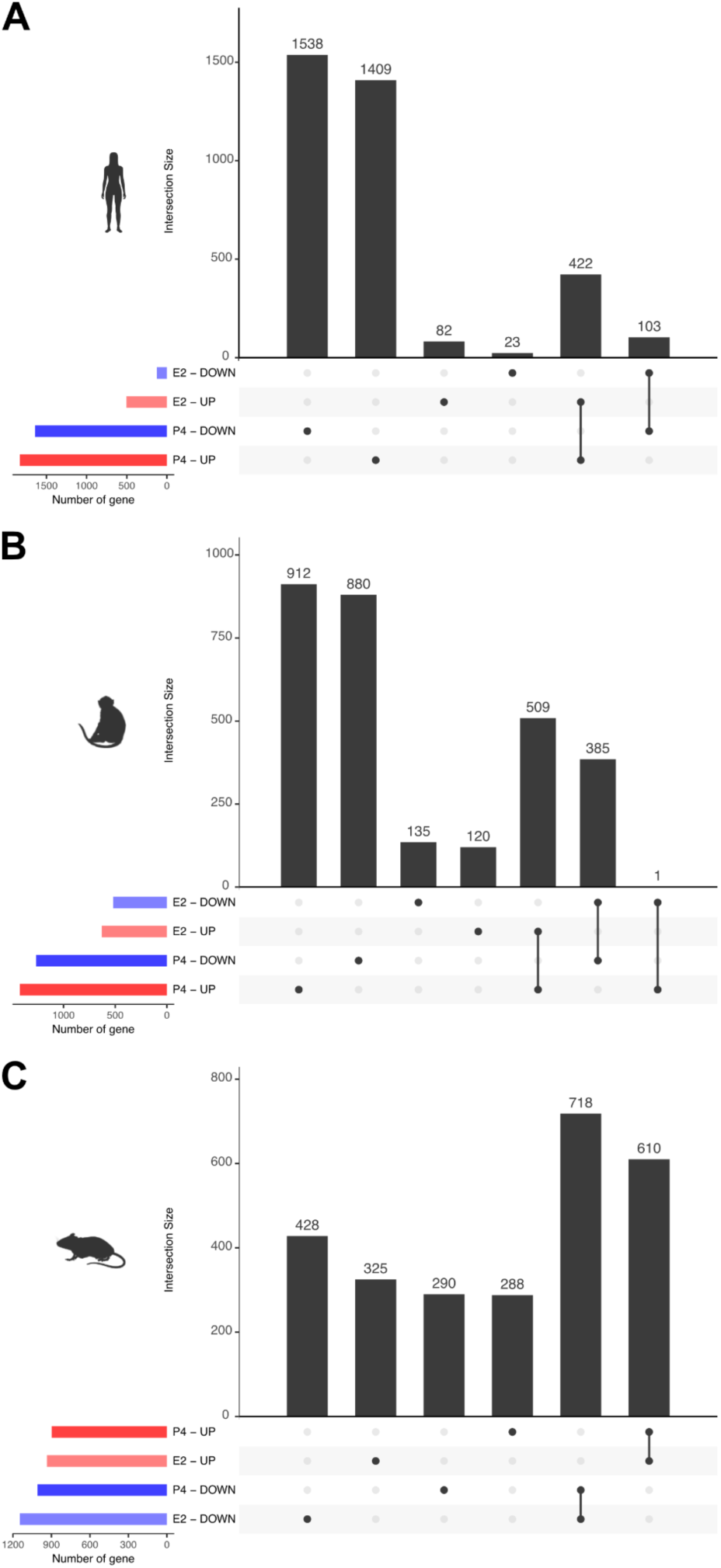
Shared DEG in response to E2 or E2+P4 treatment in each species. Upset plot representing the number of shared genes up- or down-regulated in response to E2 and E2+P4 in **(A)** human, **(B)** macaque and **(C)** mouse.

**Figure S6.**
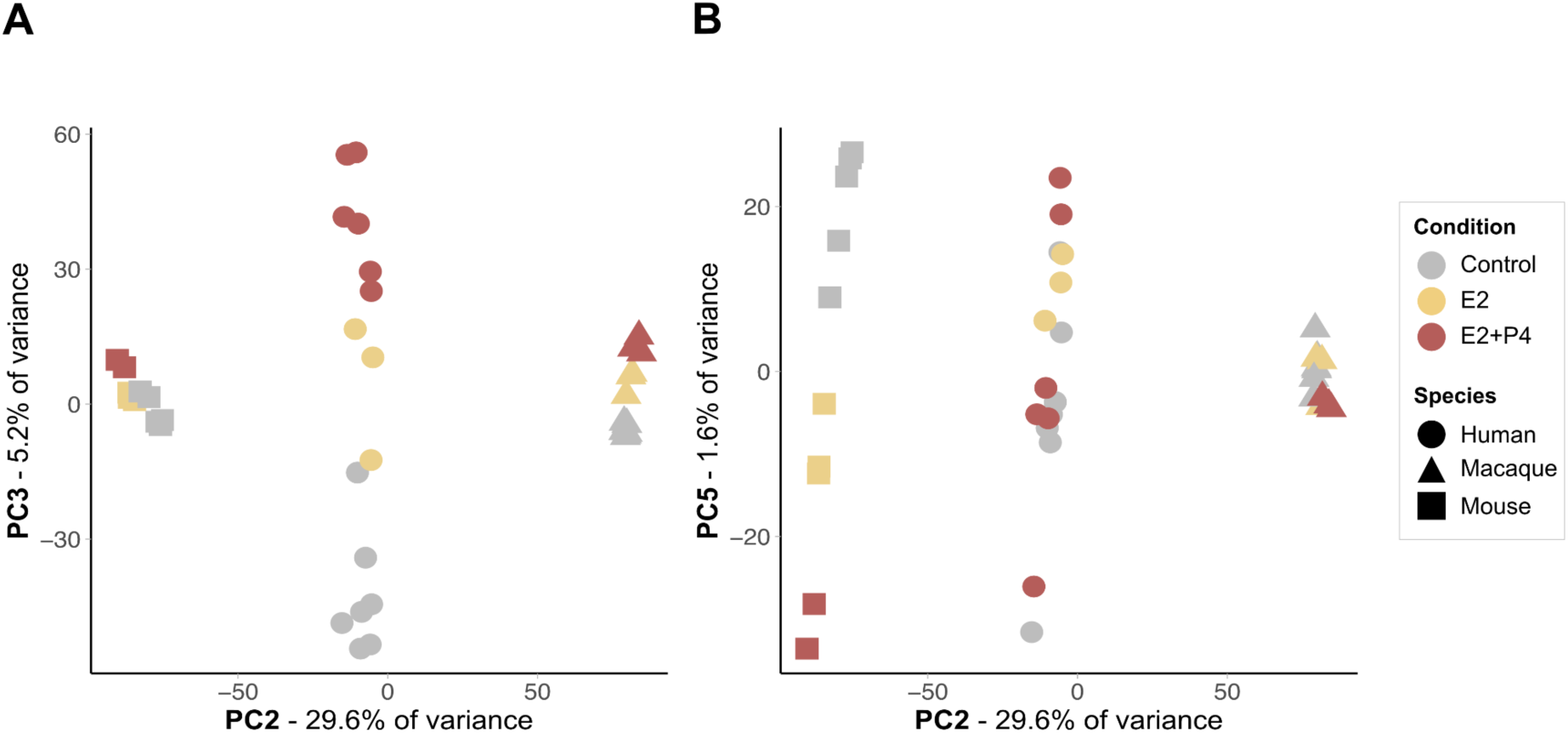
Principal component analysis of organoids bulk RNAseq across species. PCA of gene expression across 15,015 one-to-one orthologues between all three species showing (**A**) PC2 *vs* PC3, which correlates with hormonal treatment in human and macaque, and (**B**) PC2 *vs* PC5, which correlates with hormonal treatment in mouse.

**Figure S7.**
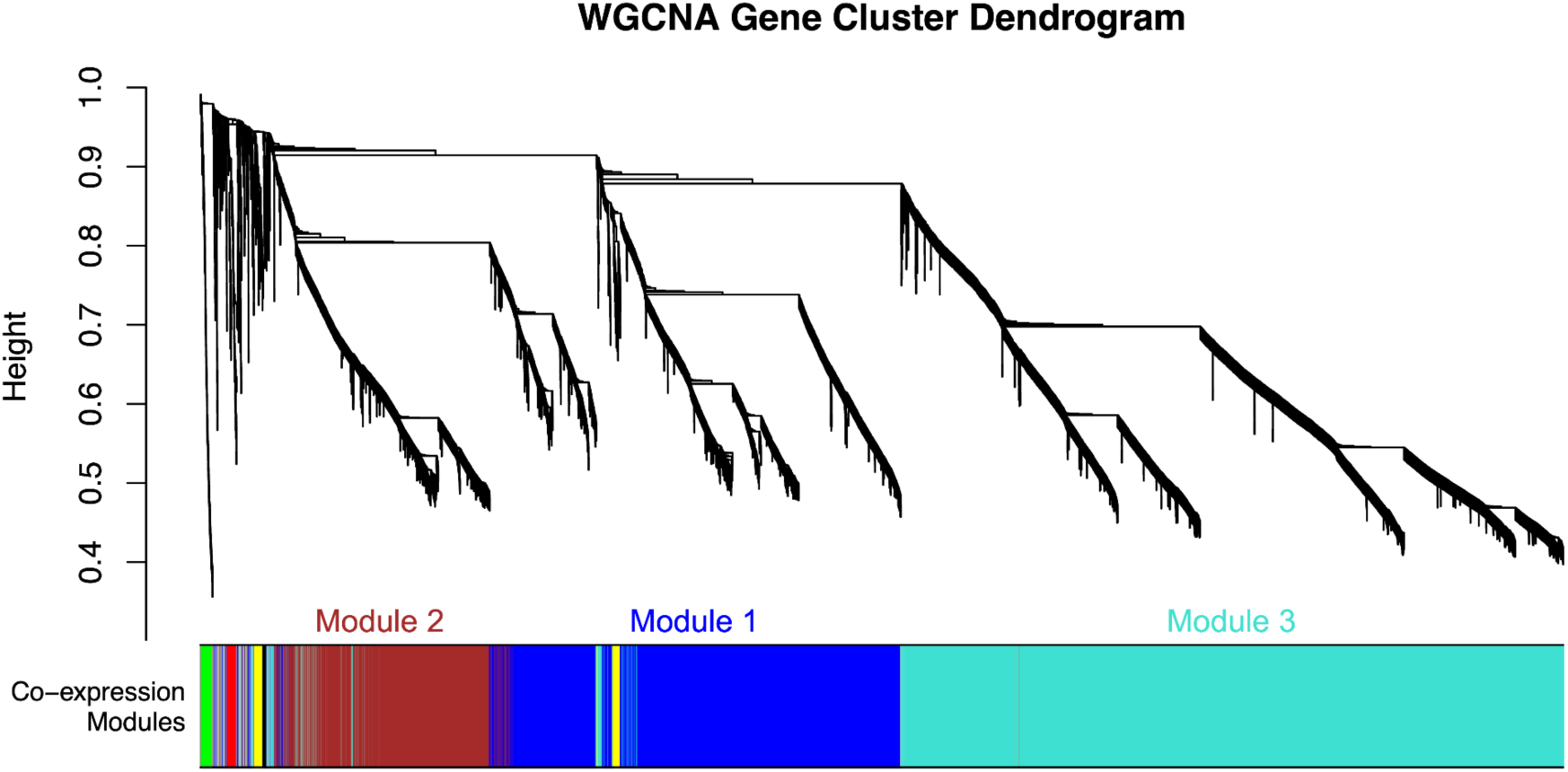
WGCNA dendrogram and gene modules. Clustering dendrogram of gene co-expression measures produced by WGCNA (Weighted Gene Correlation Network Analysis), with dissimilarity based on topological overlap.

**Figure S8.**
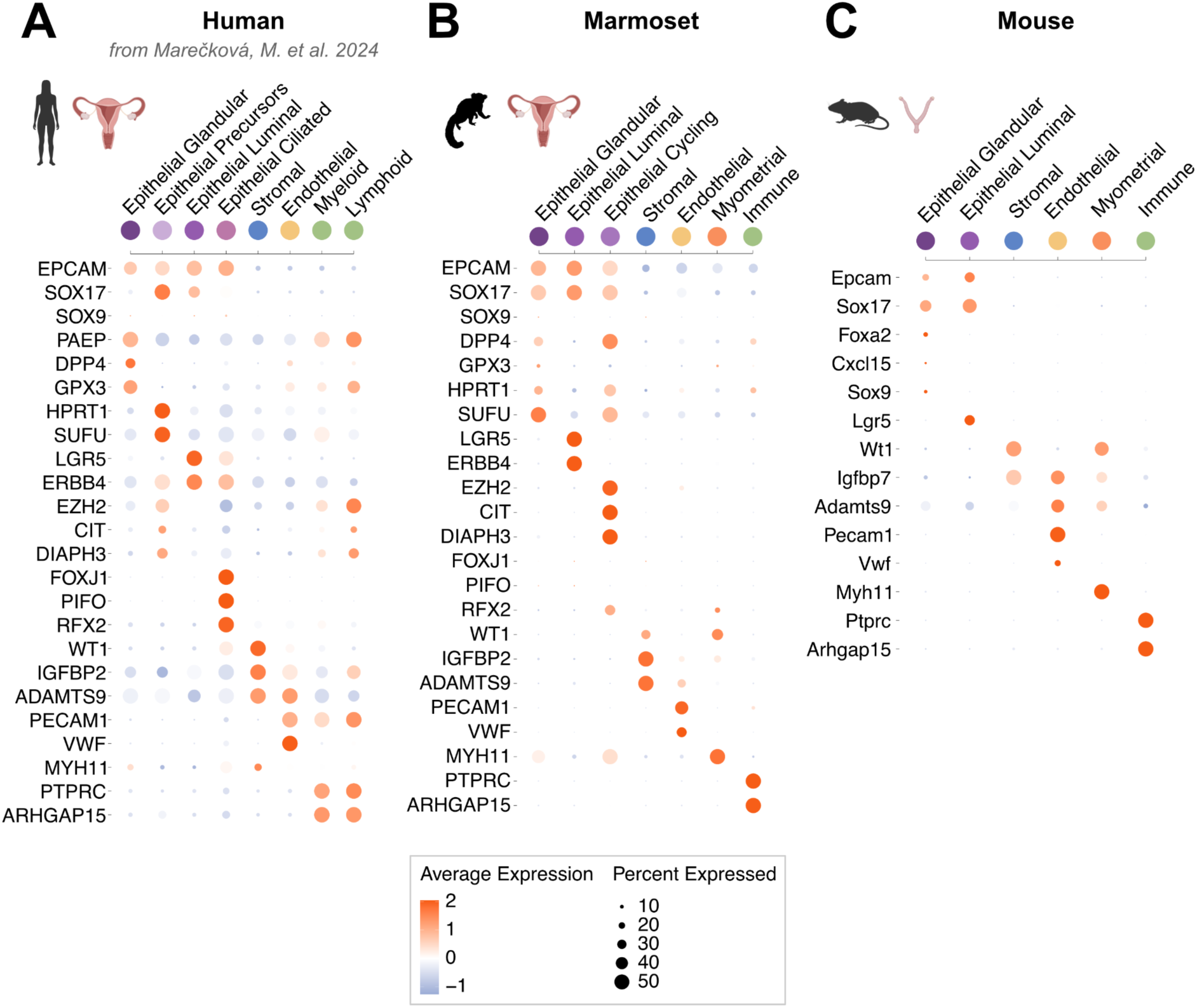
Marker genes for endometrial cell types annotation from single-nuclei RNA-seq data. **(A)** Human, (**B**) marmoset and (**C**) mouse expression of representative marker genes in the different cell types identified by single-nuclei transcriptome sequencing from Fig. 4A. Dot size indicates the fraction of cells expressing a gene, and color indicates the scaled average expression by cell (z-score) (Methods).

**Figure S9.**
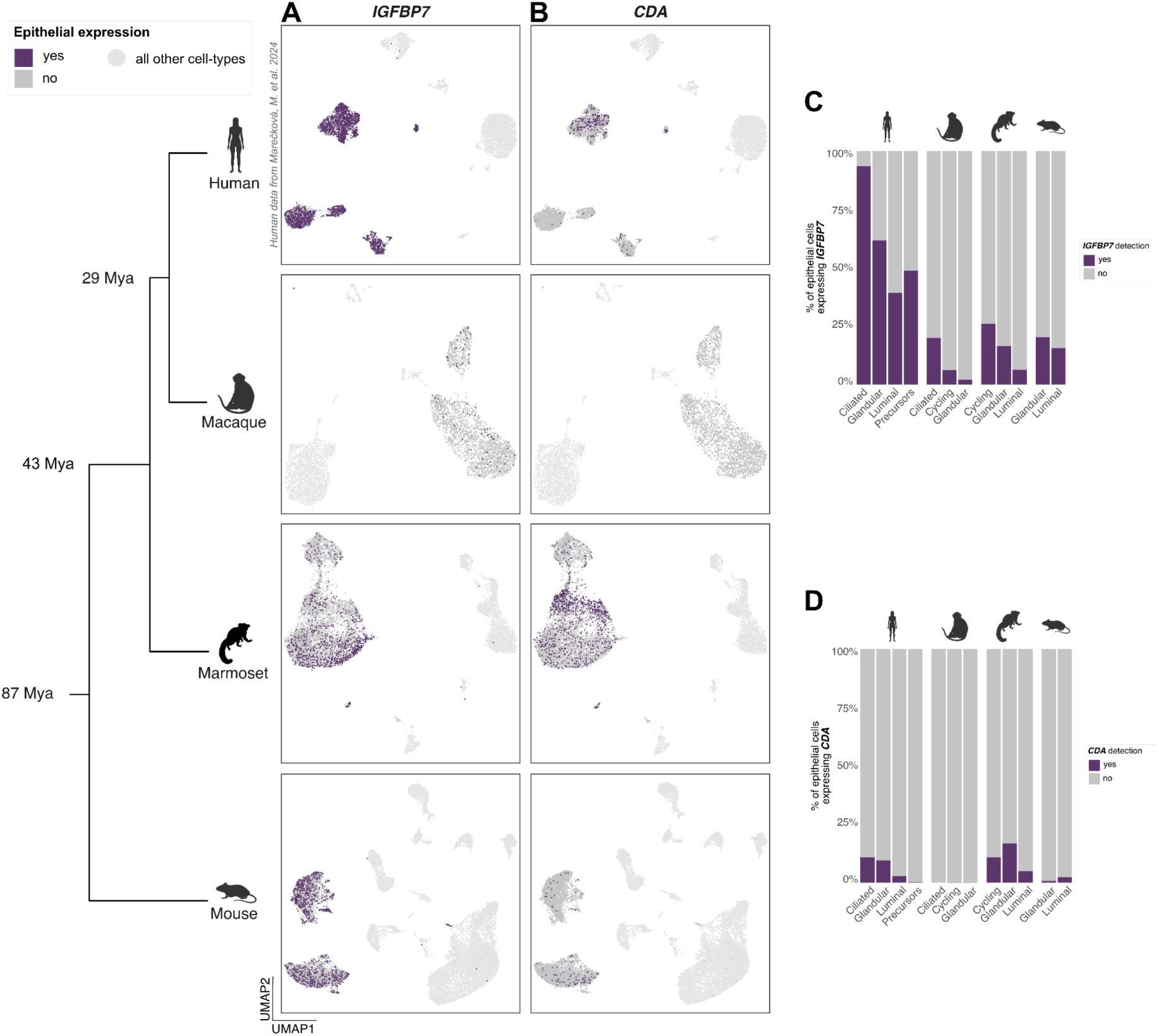
*In vivo* expression of *IGFBP7* and *CDA* expression in epithelial cells during the window of implantation in human, macaque, marmoset and mouse. (**A-B**) Expression of *IGFBP7* (B) and *CDA* (C) in epithelial cells in each species. Epithelial cells where expression was detected (gene count ≥ 1) are colored in purple. Epithelial cells where no expression was detected are colored in grey. Other cell types are colored in light grey. (**C-D**) Fractions of cells expressing *IGFBP7* (**C**) and *CDA* (**D**) across epithelial cell subtypes in each species.

## Supplementary tables

The supplementary Excel file is archived on Zenodo: https://doi.org/10.5281/zenodo.14710581

**Table S1 : DEGs common to all species after each treatment** (supplementary Excel file)

**Table S2 : Shiny GO output for DEGs common to all species after each treatment** (supplementary Excel file)

**Table S3: Manually curated list of maternal implantation genes** (supplementary Excel file)

**Table S4: List of genes co-expressed with LIF** (supplementary Excel file)

**Table S5:** Growth medium composition for mouse endometrial organoids.

| Product | Company | Product Number | Final Concentration |
| --- | --- | --- | --- |
| Advanced DMEM/F12 | Gibco | 12634-010 | 1X |
| N2 supplement | ThermoFischer | 17502048 | 1% |
| B27 supplement | ThermoFischer | 17504044 | 2% |
| Glutamax supplement | ThermoFischer | 35050038 | 2 mM |
| Penicillin/streptomycin | Gibco | 15140-148 | 1% |
| Insulin/transferrin/selenium | ThermoFischer | 41400045 | 1% |
| Nicotinamide | Sigma Aldrich | 72340 | 1 mM |
| Recombinant mouse EGF | R&D Systems | 2028 – EG | 50 ng/ml |
| Recombinant human Noggin | R&D Systems | 6057 NG | 100 ng/ml |
| Recombinant human Rspodin-1 | R&D Systems | 4645-RS | 140 ng/ml |
| Recombinant human FGF-10 | Peptotech | 100-26 | 50 ng/ml |
| Recombinant human FGF2 | R&D Systems | 233-FB | 2 ng/ml |
| Recombinant mouse WNT3A | Peptotech | 31S-20 | 250 ng/ml |
| Rock Inhibitor Y-27632* | Merk Millipore | SCM075 | 10 uM |
| ALK-4, -5, -7 inhibitor, A8301 | Sigma Aldrich | SMLO788 | 0.5 uM |

\*Only at seeding and for the first refreshment of the medium at passage 0 (i.e. for 4 days) and at each passage for 48 hours.

**Table S6:** Growth medium composition for macaque endometrial organoids.

| Product | Company | Product Number | Final Concentration |
| --- | --- | --- | --- |
| Advanced DMEM/F12 | Gibco | 12634-010 | 1X |
| N2 supplement | ThermoFischer | 17502048 | 1% |
| B27 supplement | ThermoFischer | 17504044 | 2% |
| Glutamax supplement | ThermoFischer | 35050038 | 2 mM |
| Penicillin/streptomycin | Gibco | 15140-148 | 1% |
| Insulin/transferrin/selenium | ThermoFischer | 41400045 | 1% |
| Nicotinamide | Sigma Aldrich | 72340 | 1 mM |
| N-Acetyl-L-cysteine | Sigma Aldrich | A7250 | 1.25 mM |
| Recombinant human EGF | R&D Systems | 2028 – EG | 50 ng/ml |
| Recombinant human Noggin | R&D Systems | 6057 NG | 100 ng/ml |
| Recombinant human Rspodin-1 | R&D Systems | 4645-RS | 70 ng/ml |
| Recombinant human FGF-10 | Peptotech | 100-26 | 50 ng/ml |
| Recombinant human FGF2 | R&D Systems | 233-FB | 2 ng/ml |
| Recombinant mouse WNT3A | Peptotech | 31S-20 | 250 ng/ml |
| Rock Inhibitor Y-27632* | Merk Millipore | SCM075 | 10 uM |
| ALK-4, -5, -7 inhibitor, A8301 | Sigma Aldrich | SMLO788 | 0.5 uM |
| SB202190 (p38i) | Sigma Aldrich | S7067 | 10 uM |

\*Only at seeding and for the first refreshment of the medium at passage 0 (i.e. for 4 days) and at each passage for 48 hours.

**Table S7:**
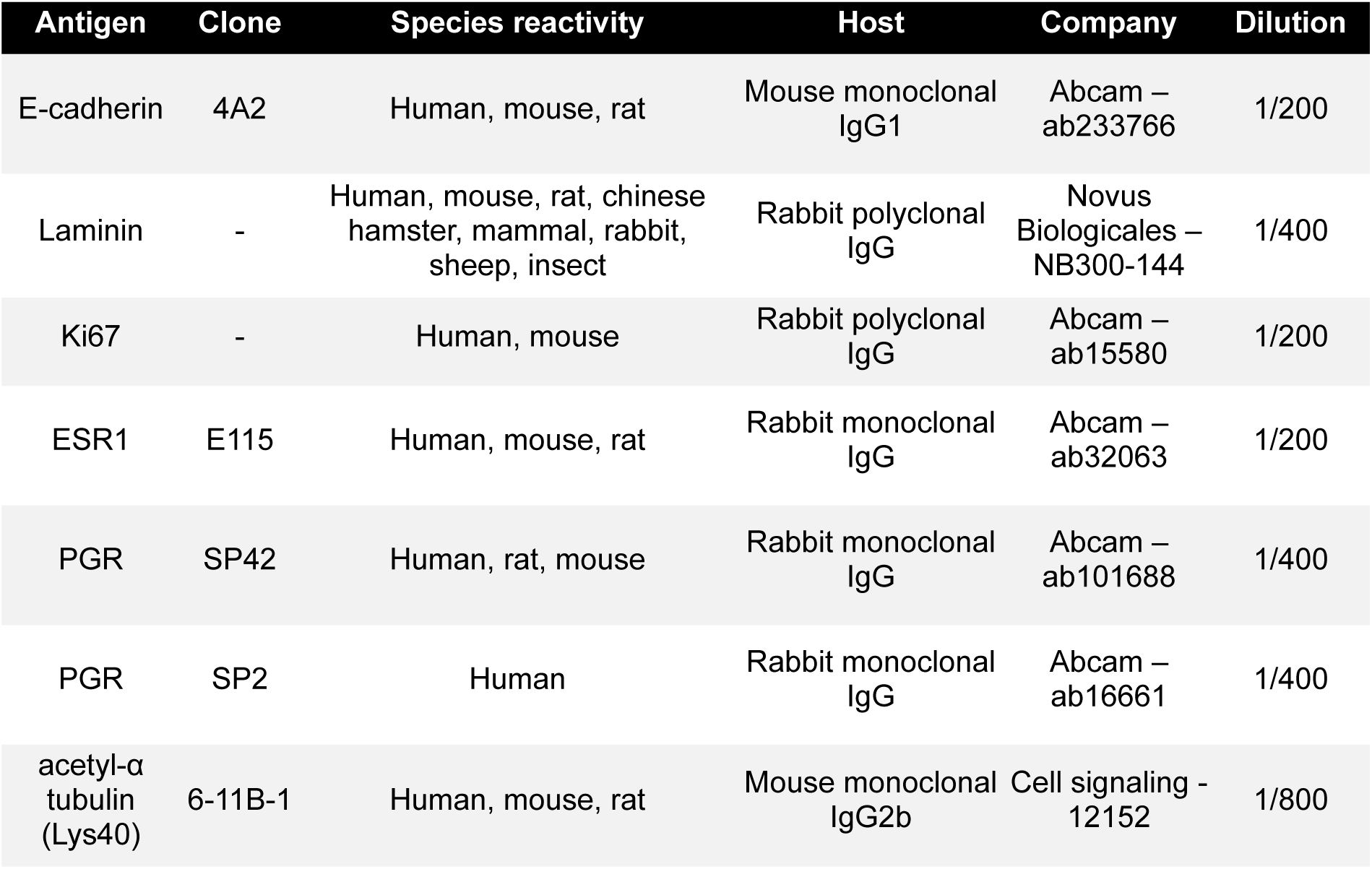
Antibodies for immunohistochemical and immunofluorescence staining.

| Antigen | Clone | Species reactivity | Host | Company | Dilution |
| --- | --- | --- | --- | --- | --- |
| E-cadherin | 4A2 | Human, mouse, rat | Mouse monoclonal IgG1 | Abcam – ab233766 | 1/200 |
| Laminin | - | Human, mouse, rat, chinese hamster, mammal, rabbit, sheep, insect | Rabbit polyclonal IgG | Novus Biologicales – NB300-144 | 1/400 |
| Ki67 | - | Human, mouse | Rabbit polyclonal IgG | Abcam – ab15580 | 1/200 |
| ESR1 | E115 | Human, mouse, rat | Rabbit monoclonal IgG | Abcam – ab32063 | 1/200 |
| PGR | SP42 | Human, rat, mouse | Rabbit monoclonal IgG | Abcam – ab101688 | 1/400 |
| PGR | SP2 | Human | Rabbit monoclonal IgG | Abcam – ab16661 | 1/400 |
| acetyl-α tubulin (Lys40) | 6-11B-1 | Human, mouse, rat | Mouse monoclonal IgG2b | Cell signaling - 12152 | 1/800 |

**Table S8 : Endometrial epithelial organoid bulk RNA-seq quality metrics** (supplementary Excel file)

**Table S9 : Endometrial snRNA-seq quality metrics** (supplementary Excel file)

**Table S10 : Marker genes for macaque single-nuclei clusters** (supplementary Excel file)

**Table S11 : Marker genes for marmoset single-nuclei clusters** (supplementary Excel file)

**Table S12 : Marker genes for pseudopregnant mouse single-nuclei clusters** (supplementary Excel file)

